# Condensation of LINE-1 is required for retrotransposition

**DOI:** 10.1101/2022.04.11.487880

**Authors:** Srinjoy Sil, Jef D Boeke, Liam J Holt

## Abstract

LINE-1 (L1) is the only autonomously active retrotransposon in the human genome, and accounts for 17% of the human genome. The L1 mRNA encodes two proteins, ORF1p and ORF2p. ORF1p is a homotrimeric RNA-binding protein that plays a critical role in assembling functional L1 ribonucleoprotein (RNP) complexes. Here we show that condensation of ORF1p is required for L1 retrotransposition. Using a combination of biochemical reconstitution and live-cell imaging, we demonstrate that RNA binding, electrostatic interactions, and trimer conformational dynamics together tune the properties of ORF1p assemblies to allow for efficient L1 condensate formation in cells. Furthermore, we directly relate the dynamics of ORF1p assembly to the ability to complete the entire retrotransposon life-cycle. Mutations that prevented ORF1 condensation led to loss of retrotransposition activity, while orthogonal restoration of coiled-coil conformational flexibility rescued both condensation and retrotransposition. Based on these observations, we propose that ORF1p oligomerization on L1 RNA drives the formation of a dynamic L1 condensate that is essential for retrotransposition.

## Introduction

Retrotransposons are genetic elements that are able to replicate themselves within a host genome in a process known as retrotransposition, a “copy and paste” mechanism that utilizes an RNA intermediate. The Long Interspersed Nuclear Element 1 (LINE-1 or L1) family of retrotransposons comprises 17% of the human genome by sequence and is the only autonomously active retrotransposon in the human genome, encoding proteins necessary for its own transposition that also drive propagation of non-autonomous retrotransposons and processed pseudogenes, which make up an additional 21% of the human genome (Lander et al. 2001; Jurka 1997; Dewannieux, Esnault, and Heidmann 2003; Haig H. Kazazian Jr and Moran 2017). While the majority of genomic human L1s have undergone truncations and mutations that have rendered them inactive, full-length, retrotransposition-competent L1s are 6 kilobases (kb) in length and have a 5’ untranslated region (UTR) containing a bidirectional promoter, two open reading frames (ORFs), ORF1 and ORF2, separated by a short inter-ORF linker, and a 3’ UTR with a weak polyadenylation signal (Figure 1A) (Speek 2001; Swergold 1990; Dombroski et al. 1991; Doucet et al. 2015; Burns and Boeke 2012).

**Figure 1:**
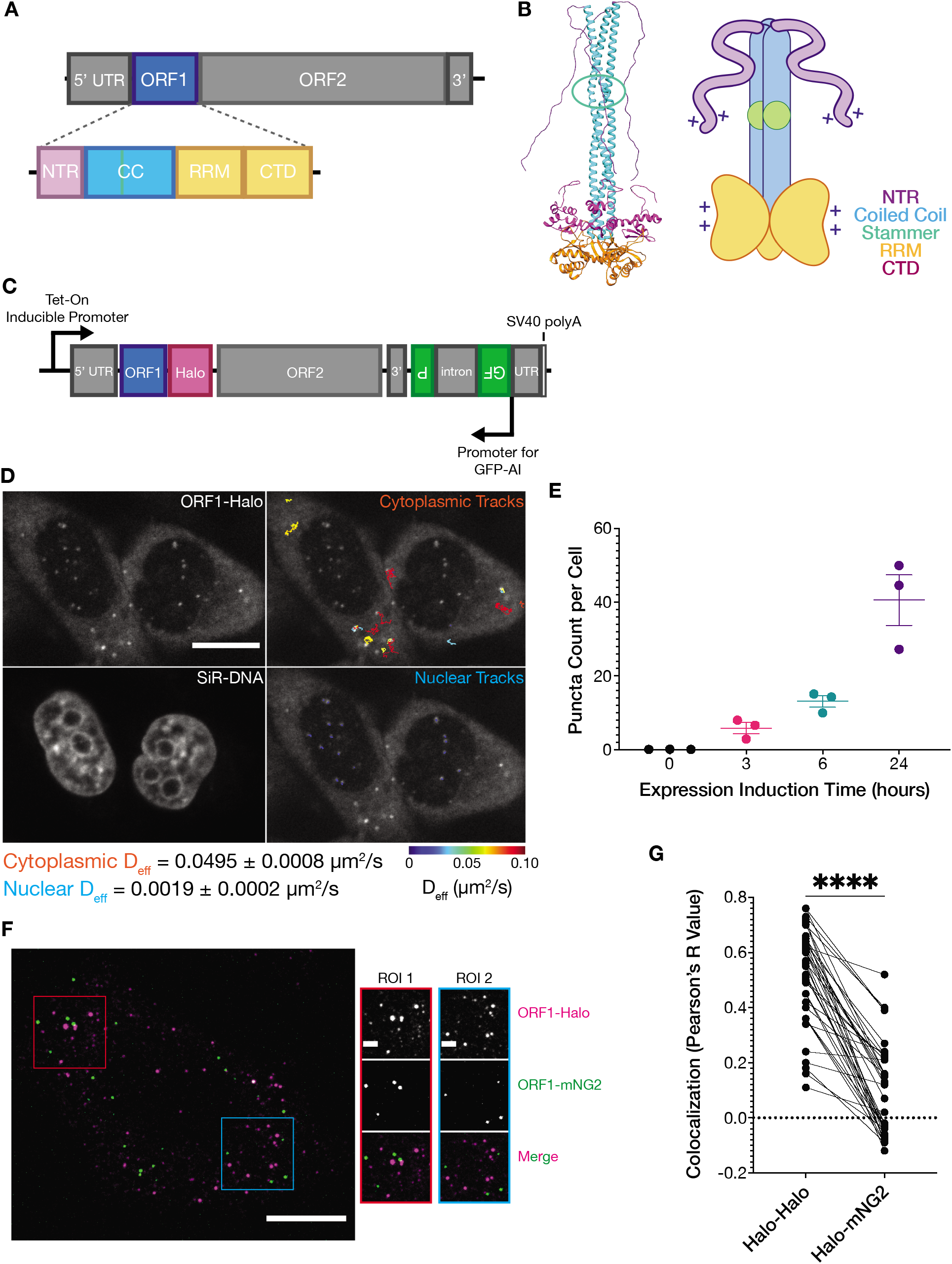
ORF1p forms stereotyped diffusive puncta in live cells that do not readily mix. **(A)** Schematic of a full-length endogenous L1 element, with a detailed view of the domains of ORF1p. **(B)** A structural model of an ORF1p trimer. A composite model generated by superimposing the coiled-coil structure (Protein Data Bank (PDB) entry 6FIA) on the RRM and CTD structure (PDB entry 2YKO). An extended conformation of the disordered NTR is modeled in. The flexibility-conferring stammer motif in the coiled coil is circled in green. A cartoon model highlighting the motifs of interest is presented on the right. **(C)** Schematic of the modified L1RP element used for cellular expression. A Tet-On CMV promoter drives expression of the full-length L1, which also contains a C-terminal HaloTag7 (Halo) on ORF1p and a GFP-AI retrotransposition reporter in its 3’ UTR. **(D)** ORF1 puncta diffuse much more slowly in the nucleus compared to the cytoplasm. A representative confocal micrograph at a single Z position shows ORF1 puncta in live HeLa cells after 24 hours of expression (top left) and corresponding nuclear staining (bottom left). Particle tracks generated from 10 seconds (100 frames) of imaging are shown in the cytoplasm (top right) and nucleus (bottom right) and are colored from blue to red, with blue indicating low effective diffusion and red representing high. Reported effective diffusion (D_eff_) is the median and SEM D_eff_ of 20 fields of view that each contain more than 5 puncta-containing HeLa cells following 24 hours of L1 expression. Scale bar = 5 μm. **(E)** The number of ORF1p condensates in cells increases with increased expression induction time. The average number of ORF1p puncta per cell was quantified after different L1 expression induction times. Each point represents one biological replicate of induction and is the average of at least 75 cells. The mean and SEM of 3 biological replicates are shown. **(F)** ORF1p from co-expressed L1s predominantly condense separately in singly-labeled foci. Representative image of a fixed HeLa cell expressing ORF1-Halo (magenta) and ORF1-mNG2 (green) off of two separate L1 expression constructs for 5 hours. Red and blue squares represent representative ROIs used for analysis of colocalization in (G). Scale bars = 10 μm (left) and 2 μm (ROIs). **(G)** ORF1p signal from co-expressed L1s colocalizes significantly less than a colocalization control. The cells were stained simultaneously with two Halo ligand dyes (Halo-JF549 and Halo-JF646), giving a positive control for colocalization. Analysis of colocalization between the two Halo signals was performed and compared with the paired colocalization of Halo-JF549 and mNG2 from the same ROI. Each dot represents a single analyzed ROI with a line connecting its colocalization score for each pair of channels. N=41 ROIs. Two-tailed paired t test analysis was performed as described in the methods. ****p<0.0001. NTR=N-terminal region, CC=stammer, RRM=RNA recognition motif, CTD=C-terminal domain.

To undergo retrotransposition, the genomic L1 must first be transcribed by RNA polymerase II, polyadenylated, and exported into the cytoplasm where the two encoded proteins, ORF1 protein (ORF1p) and ORF2 protein (ORF2p), are translated. These proteins are both necessary for retrotransposition and exhibit *cis* preference, a phenomenon in which they are more likely to mobilize the mRNA from which they were expressed than a co-expressed L1 mRNA (Wei et al. 2001). While the mechanism for *cis* preference remains unclear, a prevalent model is that ORF1p and ORF2p bind to the mRNA from which they were translated cotranslationally or immediately after translation (Boeke 1997). These two proteins are translated non-stoichiometrically, with a large excess of ORF1p, and co-assemble with L1 mRNA to form the L1 ribonucleoprotein (RNP), which is the functional unit of L1 (Hohjoh and Singer 1996; Taylor et al. 2013). ORF2p has at least two crucial activities required for retrotransposition, the reverse transcriptase (RT) required to produce dsDNA from the RNA template ((Feng et al. 1996; Cost et al. 2002; Mathias et al. 1991)) and the endonuclease (EN), which defines the target site in genomic DNA ((Feng et al. 1996; Cost et al. 2002; Mathias et al. 1991)). Once assembled, the L1 RNP must translocate to the nucleus where ORF2p uses its endonuclease activity to create a single-stranded nick in the DNA, and reverse transcribes the L1 mRNA into DNA. The ORF2p-encoded reverse transcriptase uses the free 3’-hydroxyl of the nicked DNA as a primer, directly synthesizing the L1 complementary DNA (cDNA) into the host genome in a mechanism called target-primed reverse transcription (TPRT) (Feng et al. 1996; Cost et al. 2002; Mathias et al. 1991).

ORF1p is a 40-kDa nucleic acid binding protein that assembles into a homotrimer, and has nucleic acid binding activity that is necessary for L1 retrotransposition (S. L. Martin and Bushman 2001; Sandra L. Martin et al. 2005; Kulpa and Moran 2005). ORF1p contains three structured domains, a coiled coil (CC) that mediates trimerization, an RNA recognition motif (RRM), and a C-terminal domain (CTD) that cooperates with the RRM to bind nucleic acids (Figure 1A-B) (Januszyk et al. 2007; Khazina and Weichenrieder 2009; Khazina et al. 2011). The N-terminal region (NTR) is the first 52 residues of ORF1p. This region is unstructured and contains phosphorylation sites and a basic charged patch that are necessary for retrotransposition (Cook, Jones, and Furano 2015; Khazina and Weichenrieder 2018; Adney et al. 2019).

Many RNA-binding proteins participate in the formation of membraneless organelles, such as nucleoli, stress granules (SGs), RNA processing bodies, and mRNA transport granules (Brangwynne et al. 2009; Brangwynne, Mitchison, and Hyman 2011; Feric et al. 2016; H. Zhang et al. 2015; Wheeler et al. 2016; Lee et al. 2020). These biomolecular condensates demix from the surrounding cytoplasm or nucleoplasm and serve specific functions by selectively including proteins and nucleic acids based on their biochemical properties (Hyman, Weber, and Jülicher 2014). The process by which these membraneless compartments form is known as phase separation, or biomolecular condensation, and requires the constituent molecules to exceed a critical concentration to undergo nucleation, seeding an assembly that can then further grow and undergo fusion events (Zwicker et al. 2014; Woodruff et al. 2017). Alternatively, condensate growth and fusion may be limited by the formation of a condensed phase with more solid-like material properties (Maharana et al. 2018) or by frustration of droplet coalescence in a complex mechanical environment (Y. Zhang et al. 2021).

Across the numerous characterized phase-separating RNA-binding proteins, shared molecular features such as multivalency, intrinsically disordered domains, and structured RNA binding have been shown to be important for condensation (Sanders et al. 2020; P. Yang et al. 2020; Molliex et al. 2015). Given that ORF1p exhibits all three of these molecular features of phase-separating proteins, we hypothesized that ORF1p undergoes condensation to carry out its roles in L1 RNP formation and chaperoning of L1 machinery. Recent work has confirmed that purified ORF1p is able to form a liquid-like condensed phase *in vitro* and that a truncated protein containing only the NTR and CC is sufficient for phase separation (Newton et al. 2021). Here we demonstrate that ORF1p expressed from a full-length active L1 element rapidly forms cytoplasmic condensates in cells. Structured RNA binding, disordered domain charge patches, and trimer dynamics are all necessary for cellular condensation. Biochemical reconstitution experiments reveal that the physical properties of the reconstituted ORF1p condensates, rather than their *in vitro* size, predict their propensity to assemble in cells. ORF1p condensation *in vivo* correlates with measured L1 retrotransposition activity, and orthogonal restoration of trimer dynamics rescues both condensation and retrotransposition. Together these results indicate that condensation is necessary for L1 retrotransposition. We propose that the biochemical properties of ORF1p are tuned to efficiently nucleate condensation cotranslationally, while also allowing for dynamic interactions between fully assembled L1 RNPs and host proteins and nucleic acids.

## Results

### ORF1p forms stereotyped foci in live cells

Previous studies focusing on the ORF1 protein in mammalian cells and tissues have reported a variety of localizations (Doucet, Basyuk, and Gilbert 2016; Goodier et al. 2004, 2007; Sharma et al. 2016; Rodić et al. 2014; Pereira et al. 2018; Mita et al. 2018). We designed an inducible, active, endogenous-like L1 expression construct in which ORF1p was fused at its C-terminus to HaloTag7 (ORF1-Halo) (Figure 1C). This approach enabled the use of fluorescence microscopy to investigate the behavior of the protein in live mammalian cells. The construct also contained a well-characterized GFP-AI retrotransposition reporter in its 3’ UTR (Ostertag et al. 2000; An et al. 2011; Mita et al. 2018). The GFP-AI cassette contains the EGFP coding sequence interrupted by the γ-globin intron. This intron is in the opposite orientation as the coding sequence and disrupts the GFP open-reading frame. A CMV promoter and a thymidine kinase (TK) poly(A) signal flank the EGFP sequence, and the entire cassette is oriented antisense to the L1 sequence. When the L1 is transcribed, the γ-globin intron is removed by splicing, and successful retrotransposition of this spliced L1 mRNA construct allows for the subsequent expression of the uninterrupted EGFP coding sequence from the novel insertion site. Using this reporter enables us to associate changes in ORF1p behavior with effects on L1 retrotransposition activity. Thus, we are able to directly relate the dynamics of ORF1p assembly to the ability to complete the entire retrotransposon life-cycle.

Upon a 6-hour induction of L1 expression using the Tet-On system in HeLa cells, we noted the formation of bright, punctate, and highly uniform ORF1-Halo structures against a weaker background of diffuse Halo signal (Figure 1D, Figure 1-Supp 1A). These foci were absent without induction, appeared as early as three hours following induction with doxycycline, and increased in number, but not size, with longer induction times (Figure 1E, Figure 1-Supp 1A-B). The ORF1 puncta that we describe in live cells are smaller and more uniform than previously visualized ORF1 assemblies in fixed cells (Goodier et al. 2007; Taylor et al. 2013; Mita et al. 2018). We observe similar stress-granule-like ORF1 condensates after inducing expression of our L1 construct for 72 hours (Figure 1-Supp 1C), suggesting that these larger assemblies are a result of higher ORF1p expression levels while the uniform puncta seen after 6 hours may be more representative of ORF1p behavior at physiological expression levels. Although the diffuse ORF1 signal remained predominantly cytoplasmic, nuclear ORF1 foci began to appear at longer induction times. Nuclear ORF1p puncta had similar morphology and fluorescence intensity but very different diffusivity compared to cytoplasmic puncta (Figure 1D, Figure 1-Supp 1D). Cytoplasmic ORF1 puncta had readily observable diffusion at millisecond timescales, while nuclear puncta were virtually immobile at similar imaging speeds (Supp Movie 1). In summary, upon induction of L1 expression, ORF1p quickly forms stereotyped puncta that diffuse rapidly in the cytoplasm, while nuclear ORF1 puncta take longer to appear and exhibit starkly reduced diffusivity.

ORF1p and ORF2p exhibit preferential binding to the L1 mRNA from which they were translated, thus promoting transposition of intact and active L1s rather than mobilizing other L1-derived RNAs (Boeke 1997); this phenomenon is known as *cis* preference, but its molecular mechanism remains unclear. There is functional evidence for the *cis* preference of ORF1p, as expression of an intact L1 minimally complemented the retrotransposition of a co-expressed marked L1 encoding an ORF1 with an RNA-binding deficiency (Wei et al. 2001; Kulpa and Moran 2006). Additionally, the L1s that were identified in cases of novel insertional mutagenesis were always derived from intact full-length L1s, suggesting that active L1s are far more likely to drive *cis* retrotransposition than they are to mobilize the much more common mutated or truncated L1s (H. H. Kazazian Jr et al. 1988; Woods-Samuels et al. 1989; Dombroski et al. 1991; Holmes et al. 1994; Moran et al. 1996; Brouha et al. 2003; Boeke 1997). We therefore wondered whether the ORF1p assemblies that we observed would form cotranslationally and stay separate, or conversely, could mix with each other, either directly through fusion events or indirectly through protein exchange with the surrounding cytoplasm. To answer this question, we simultaneously expressed two separate L1 elements differing only in the type of tag on the ORF1p (HaloTag7 or mNeonGreen2 (mNG2)) in the same cell. We used the JF549 Halo ligand and 561-nm laser illumination to visualize ORF1-Halo (Figure 1F, magenta) and a 488-nm laser to visualize ORF1-mNG2 (Figure 1F, green). We almost always found singly-labeled puncta and only rarely found dual-labeled puncta (Supp Movie 2). As a positive control for colocalization, cells co-expressing ORF1-Halo and ORF1-mNG2 were simultaneously stained with JF646 Halo ligand in addition to JF549 prior to fixation, and the JF646 signal was visualized using 640-nm laser illumination. Since the two Halo ligand signals should strongly colocalize, we used their colocalization as a comparison for the colocalization of the Halo and mNG2 signals in the same region of interest (ROI) and found that the ORF1-Halo and ORF1-mNG2 had significantly lower colocalization than the positive control (Figure 1G). These results suggested that ORF1p assemblies do not rapidly exchange protein with the surrounding cytoplasm or undergo frequent fusion events and are perhaps instead kinetically trapped in an assembled form cotranslationally. This observation is consistent with previous work on the functional *cis* preference of L1-encoded proteins (Wei et al. 2001; Kulpa and Moran 2006). In this way we showed that ORF1p puncta undergo minimal mixing in live cells, suggesting a role for rapid ORF1p assembly in forming *cis*-preferential L1 RNPs.

### Purified ORF1p forms liquid-like droplets and co-condenses with RNA in vitro

Next, we used *in vitro* biochemistry to determine whether a minimal reconstituted system could form condensates *in vitro.* To this end, we purified full-length ORF1p from *E. coli.* We modified previously described protocols (Carter et al. 2020) to maximize removal of protein-bound RNA (Figure 2-Supp 1, methods). We fluorescently labeled the purified protein using an amine-reactive fluorescent dye (Nanda and Lorsch 2014) and used a ratio of at least 10:1 unlabeled:labeled ORF1p to visualize protein distribution in microscopy assays. Purified ORF1p protein formed an extensive condensed phase at low micromolar protein concentrations in buffer with physiological pH and salt concentrations (Figure 2A). When the same concentration of protein was incubated in buffers with increasing salt concentrations, the mean intensity of the protein in the condensed phase, the partition coefficient of the protein, and the total condensed phase area all decreased, with no condensate formation observed above 300 mM KCl (Figure 2D, left). Decreasing protein concentration at a fixed salt concentration decreased the total condensed phase area but increased the protein partition coefficient. At 200 and 300 mM KCl, the range of protein concentrations tested spanned the phase boundary such that higher protein concentrations led to detectable condensate formation but lower protein concentrations did not. Taken together, these experiments demonstrated that purified ORF1p robustly phase separates at physiological salt concentrations. Inhibition of phase separation at higher salt concentrations suggests that electrostatic interactions play an important role in condensation.

**Figure 2:**
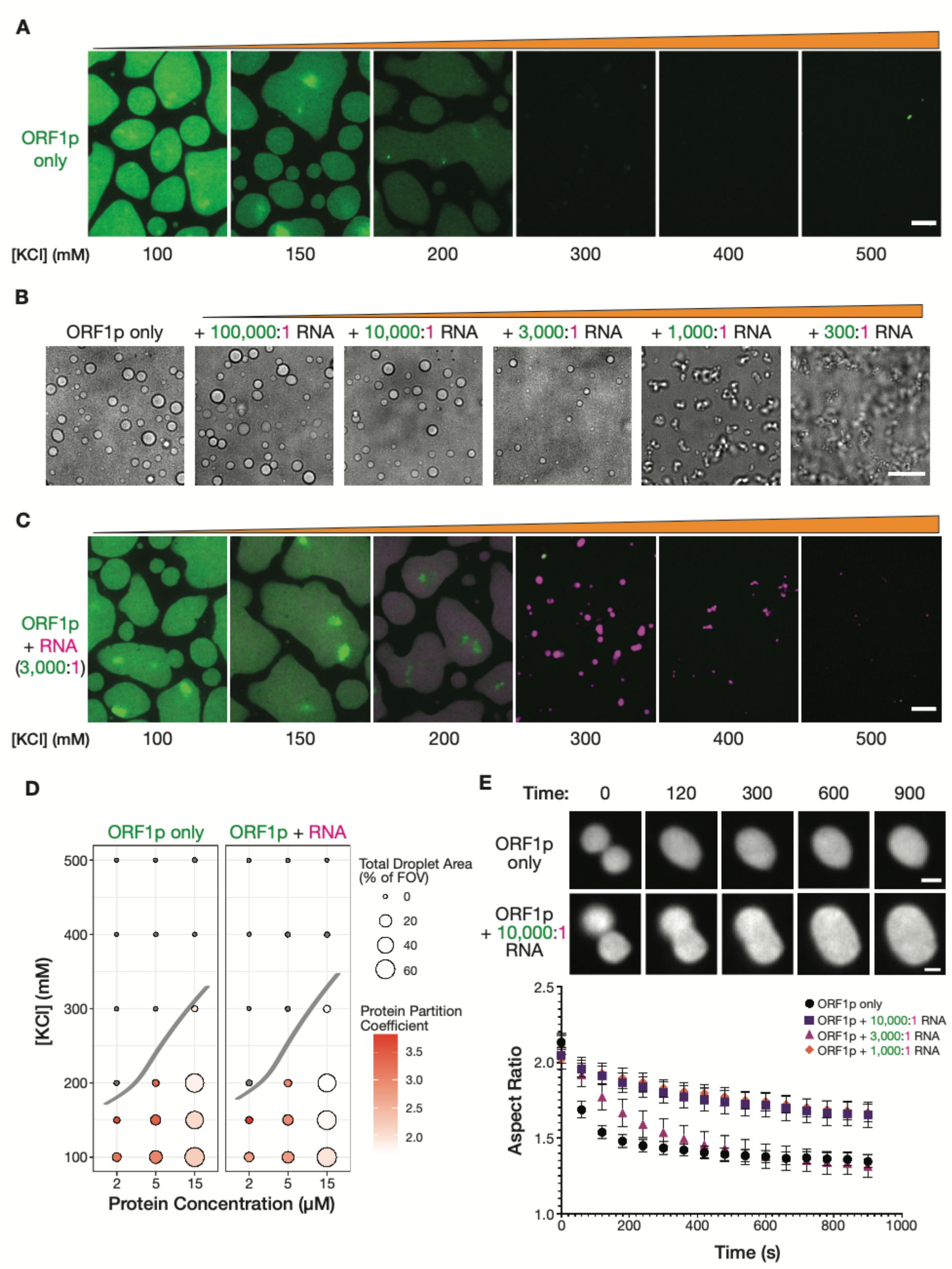
Purified ORF1p forms condensates with and without RNA, exhibiting differential condensate properties. **(A)** Purified ORF1p forms an extensive condensed phase *in vitro.* Representative images of ORF1p (green) condensed phase formation across a range of salt concentrations. 15 μM protein was used. All of the images use the same lookup tables. Scale bars = 5 μm. **(B)** Increasing RNA leads to decreasing droplet area and eventually the formation of irregular three-dimensional fibrillar structures. Representative brightfield images of ORF1p droplet morphology over a wide range of 2-kb L1 RNA stoichiometries, with increasing RNA concentration from left to right. 5 μM protein and 150 mM KCl was used in all conditions. Scale bar = 10 μm. **(C)** RNA robustly co-condenses with ORF1p *in vitro*. Representative images of ORF1p (green) condensed phase formation across a range of salt concentrations as in (A), in the presence of added labeled 2-kb L1 RNA (magenta). 15 μM protein and 5 nM RNA were used (3,000:1 protein:RNA). All of the images use the same lookup tables for each channel. Scale bars = 5 μm. **(D)** RNA addition does not strongly affect the phase diagram of ORF1p *in vitro.* A phase diagram of ORF1p with and without added RNA (3,000:1 protein:RNA). Total condensed phase area of each condition is shown by the area of the circle for each condition, and the protein partition coefficient is represented by the filling. A hand-drawn phase boundary separates conditions with appreciable condensation with those that do not. **(E)** RNA-containing ORF1p condensates have slower droplet fusion kinetics than protein-only condensates. Representative images of fusion events over 15 minutes are shown for ORF1p with and without 10,000:1 protein:RNA addition, demonstrating slower fusion with RNA. 10 μM protein and 150 mM KCl were used in all conditions. Average aspect ratios across individual fusion events in each RNA condition are plotted (Mean ± SEM) over time for 15 minutes. 10 or more fusions were analyzed per condition. Scale bars = 1 μm.

Since ORF1p has previously been described to have nucleic acid binding activity, we were curious how addition of RNA would affect ORF1p condensed phase formation. We generated a 2-kb fluorescently labeled RNA corresponding to the 5’ of the L1 mRNA for use in *in vitro* condensation assays. We used this 2-kb fragment because it was difficult to consistently generate full-length L1 RNA by *in vitro* transcription. It has been estimated that an RNA that is fully coated by ORF1p could be bound by one ORF1p trimer every 75 nucleotides (Khazina et al. 2011; Taylor et al. 2013). The full length 6-kb L1 mRNA would therefore require ~80 ORF1p trimers or 240 ORF1p molecules to fully cover the L1 mRNA, and our 2-kb RNA would require 30 trimers or 90 ORF1p molecules, resulting in a predicted ORF1p:RNA ratio of 90:1. We decided to explore the effects of a large range of RNA stoichiometries on ORF1p droplet formation at a fixed protein concentration (Figure 2B). While high ORF1p:RNA stoichiometries had minimal effects on droplet formation and morphology, we noted that the ORF1p droplets became less spherical at 1,000:1 ORF1p:RNA, with the dominant species appearing to be chains of slowly fusing droplets. At 300:1 ORF1p:RNA, the ORF1p condensed phase was primarily composed of a large network of branched fibrillar structures. The formation of these fibrillar structures at 300:1 protein:RNA was surprising, since a fully occupied 2-kb RNA is predicted to be able to accommodate 90 ORF1p molecules (Khazina et al. 2011; Taylor et al. 2013). However, our reconstitution approach mixes preformed ORF1p with RNA, which is distinct from the cotranslational assembly process that is likely to occur in cells. The kinetics of these assembly processes are very different and may lead to different material properties. Furthermore, our 2-kb RNA has distinct polymer properties compared to the full-length 6-kb L1 RNA, and we speculate that our RNA may promote fibrillarization at higher protein:RNA stoichiometries. Finally, the simple reconstituted system does not contain host factors (e.g. RNA helicases (Tauber et al. 2020; Li et al. 2013; Goodier, Cheung, and Kazazian 2012)), nor does it recapitulate the complex, crowded intracellular environment, which can have dramatic effects on phase separation (Delarue et al. 2018). Nevertheless, despite the limitations of our reconstitution approach, we showed that the physical properties of ORF1p condensates are strongly impacted by RNA abundance, with increasing RNA concentrations leading to slower droplet fusion kinetics and eventually the formation of fibrillar structures rather than droplets.

We wondered how RNA addition affected the propensity of ORF1p to phase separate in buffers with increasing salt concentrations. To enable comparison of the properties of the RNA-containing droplets with those containing protein alone, we used an intermediate RNA stoichiometry (3,000:1 ORF1p:RNA). This stoichiometry maintains a liquid-like condensed phase, thereby facilitating characterization by standard biophysical methods. When we mixed RNA with ORF1p at this stoichiometry, the labeled RNA was robustly recruited to the ORF1p condensed phase (Figure 2C). Introducing RNA into condensates at this stoichiometry did not strongly affect either the total condensate area or the protein partition coefficient compared to the corresponding condition without RNA (Figure 2D). Given our observations of solid-like condensation at lower protein:RNA stoichiometries, we wondered whether the RNA was instead altering the physical properties of the droplets.

The dynamics of liquid droplet fusion are influenced by the surface tension and viscosity of the liquid as well as the size of the droplets (Eggers, Lister, and Stone 1999). To assess the physical properties of the condensed phase, we analyzed droplet fusion events in time-lapse movies (Supp Movie 3). We analyzed each fusion event over time by fitting an ellipse to the fusion intermediate at each timepoint and calculating its aspect ratio. In a liquid-like fusion event, the aspect ratio will decrease exponentially to 1, which corresponds to a spherical fusion product. Analysis of other reconstituted protein droplets have shown that such fusions can occur on the order of a few seconds (Elbaum-Garfinkle et al. 2015). Droplets containing ORF1 protein alone exhibited slow fusion kinetics, requiring 3 minutes to reach a plateau of the aspect ratio (Figure 2E). Notably, the aspect ratio plateau was greater than 1, indicating that the fused droplets retained an ovoid shape. This could be explained by a high ratio of viscosity to surface tension, or could reflect a droplet aging effect in which the droplets become more solid-like over time (Jawerth et al. 2020). All RNA-containing ORF1p condensates fused more slowly than protein-only condensates, but surprisingly this decrease was non-monotonic with increasing RNA concentration: the addition of 10,000:1 and 1,000:1 protein:RNA resulted in fusions that plateaued at high aspect ratios, while 3,000:1 RNA condensates were able to fuse to similar final aspect ratios as the ORF1p protein alone but took more time to reach the plateau (Figure 2E). These experiments indicated that addition of RNA to the ORF1p condensed phase changes its viscosity and surface tension in a way that slows droplet fusion kinetics. Additionally, they suggested that physical properties of the ORF1p condensed phase may change depending on the ratio of ORF1p to RNA. Therefore, the physical properties of L1 condensates are likely to change during cotranslational assembly of RNPs as ORF1p timers are sequentially added to the forming RNP, increasing the relative amount of ORF1p over time.

### Key basic residue mutations alter ORF1p condensate properties in vitro and in cells

Electrostatic interactions modulated the properties of ORF1p droplets *in vitro,* implicating charged residues in ORF1p in the condensation process. We were particularly interested in a positive charge patch at the end of the N-terminal disordered region, as these types of charged motifs have been implicated in nucleic acid binding and protein-protein interactions in other contexts, for example in the disordered tails of transcription factors (Boija et al. 2018; Tóth-Petróczy et al. 2009). Furthermore, two lysines within this charge patch, K3 and K4, have been previously reported to decrease L1 retrotransposition in cells when mutated (Adney et al. 2019; Khazina et al. 2011; Khazina and Weichenrieder 2018). RNA interactions also affected the physical properties of ORF1p condensates in our *in vitro* experiments. Mutation of a central DNA-contacting arginine, R261, was previously shown to strongly decrease ORF1p RRM-mediated RNA binding *in vitro* and abrogated retrotransposition activity in cells (Khazina et al. 2011). Given these previous findings, we decided to investigate the condensation properties of K3A/K4A and R261A mutant proteins.

We purified full-length K3A/K4A and R261A mutant proteins. Both mutants were purified using the same protocol as wild-type (WT) ORF1p and were confirmed to be trimeric during size-exclusion chromatography (Figure 3-Supp 1), consistent with correct overall folding and assembly of the mutant proteins.

When reconstituted in buffer with physiological pH and salt concentration, both K3A/K4A and R261A ORF1p proteins formed condensed phases (Figure 3A). Notably, the total condensed phase area of both mutants was less than that of WT, as they formed smaller and more circular droplets. When assayed across a range of protein and salt concentrations, ORF1p K3A/K4A formed condensates in almost all of the conditions that WT did, while condensation of R261A was limited to only the conditions with higher protein concentrations and lower salt concentrations (Figure 3B). The decreased phase separation of the R261A mutant was unexpected, as we predicted that mutating a core RNA-binding residue would only affect condensation in the presence of RNA. We also noted that the protein partition coefficients of the R261A condensed phases were higher than their counterparts for WT and K3A/K4A. Taken together, these experiments showed that K3/K4 and R261 are not essential for protein condensation *in vitro*. Mutations in these residues instead decreased the total area of condensed phase generated and limited condensate formation in conditions with low protein concentrations or high salt concentrations.

**Figure 3:**
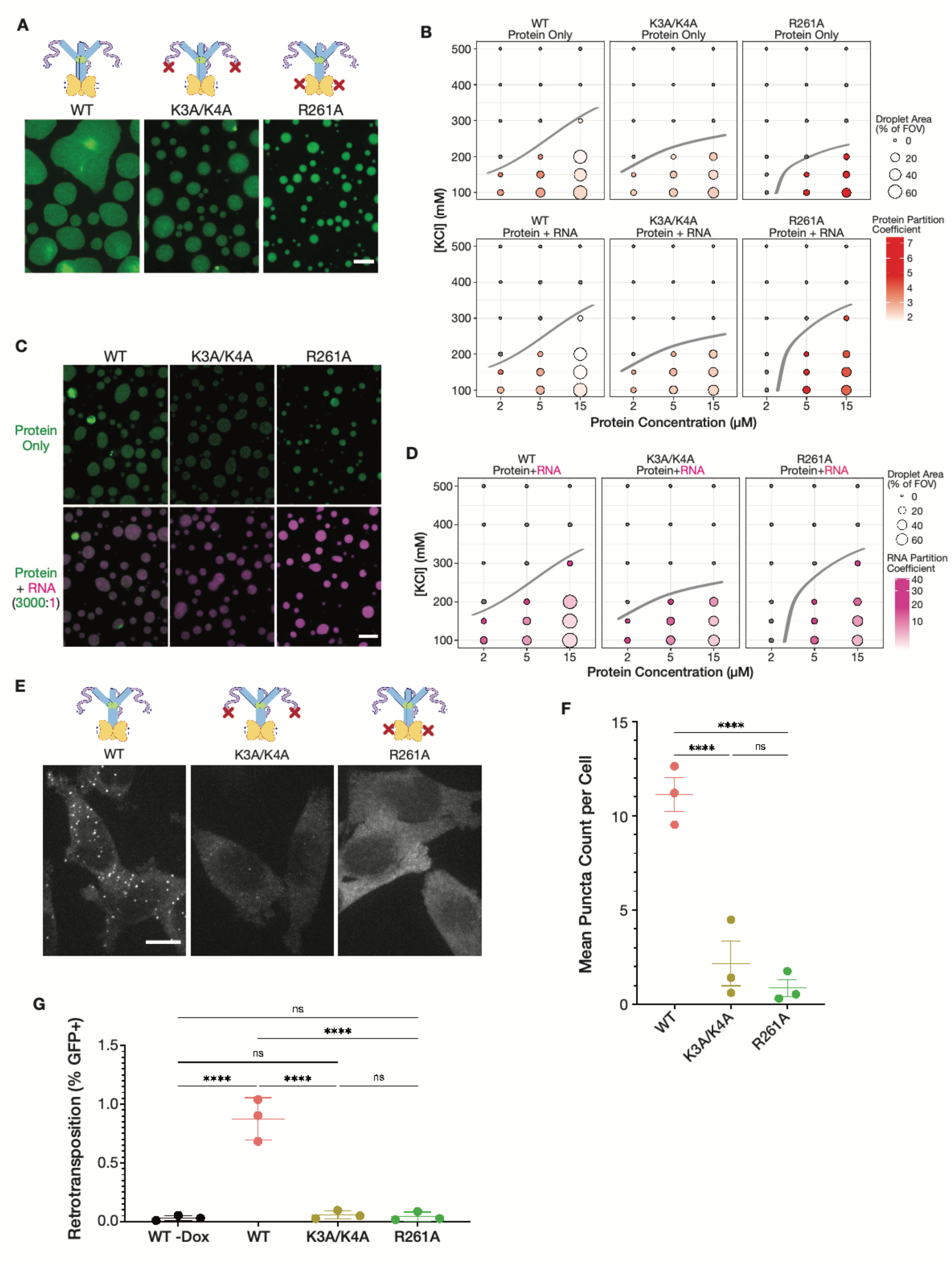
Mutations in key basic motifs modulate droplet formation *in vitro* and abrogate condensation and retrotransposition in cells. **(A)** ORF1p variants with mutations in basic motifs form condensed phases *in vitro.* Representative images of the condensed phases of WT, K3A/K4A, and R261A ORF1p. 15 μM protein and 150 mM KCl were used for all three mutants. Both mutants have reduced condensate area compared to WT. Scale bar = 5 μm. **(B)** Basic motif mutants of ORF1p have decreased condensed phase areas in conditions with high salt concentration or low protein concentration. A phase diagram of ORF1p condensation for WT, K3A/K4A, and R261A with and without the addition of 3,000:1 2-kb L1 RNA, as in Figure 2D. R261A generates an RNA-responsive condensed phase with much higher protein partition coefficients than WT or K3A/K4A. Hand-drawn phase boundaries separate conditions with appreciable condensation with those that do not. **(C)** RNA enhances R261A condensation but has minimal effect on WT and K3A/K4A. Representative images of WT, K3A/K4A, and R261A ORF1p with and without RNA (3,000:1). All condensates were generated with 5 μM protein and 100 mM KCl. Protein image LUTs are the same per mutant as in (A), and RNA LUTs are the same across mutants. Scale bar = 5 μm. **(D)** ORF1p R261A has much higher RNA partition coefficients than WT and K3A/K4A. A phase diagram showing RNA partition coefficients for WT, K3A/K4A, and R261A ORF1p with RNA (3,000:1). Handdrawn phase boundaries are included as in (B). **(E)** WT ORF1p forms cellular puncta much more robustly than K3A/K4A or R261A. Representative maximum intensity Z projections of HeLa cells expressing WT, K3A/K4A, or R261A ORF1p after 6 hours of L1 expression. All images have the same lookup tables. Scale bar = 10 μm. **(F)** Quantification of the average number of ORF1p puncta per cell after 6 hours of expression of WT, K3A/K4A, or R261A ORF1p. Each point represents one biological replicate of induction and quantification and is the average of at least 75 cells. The mean and SEM of 3 biological replicates are shown, and statistical differences between mutants were calculated using a one-way ANOVA with Tukey’s multiple comparison correction. ****p<0.0001, ns = not significant. **(G)** WT L1 retrotransposes at a cellular frequency of ~1%, while elements with ORF1 K3A/K4A and R261A have undetectable retrotransposition activity. Measured retrotransposition activity of WT, K3A/K4A, and R261A ORF1p after 72 hours of L1 expression. GFP+ cells were evaluated using FACS with a GFP+ threshold defined by WT cells without expression induction (WT -Dox). Each point is a biological replicate whose value is the average of three technical replicates, with 25,000 cells analyzed for each. The mean and SEM of 3 biological replicates are shown, and statistical differences between conditions were calculated using a one-way ANOVA with Tukey’s multiple comparison correction. ****p<0.0001, ns = not significant.

We next investigated the behavior of the mutant condensed phases in the presence of RNA. Both mutants robustly recruited the fluorescently labeled 2-kb L1 RNA into their condensates (Figure 3C). While the addition of RNA did not appreciably change the phase diagram of WT or K3A/K4A, the total area of R261A condensed phases increased modestly with the addition of RNA, and co-condensation with RNA allowed R261A condensates to stably form at higher salt concentrations than protein alone (Figure 3B-C). Additionally, the R261A condensed phases generally had higher RNA partition coefficients than WT or K3A/K4A (Figure 3D), despite the protein’s reported deficiency in binding to structured RNA (Khazina et al. 2011). These experiments demonstrated that ORF1p K3A/K4A and R261A are able to co-condense with RNA, surprisingly, with R261A condensates exhibiting increased droplet size and decreased salt sensitivity in the presence of RNA.

Given the altered condensation properties of these ORF1p mutants *in vitro,* we next sought to characterize their condensation in mammalian cells. Upon expressing WT, K3A/K4A, and R261A ORF1p in the context of a full-length L1 element for 6 hours in HeLa cells, we observed a stark reduction in puncta formation of the ORF1p mutants compared to WT. ORF1p K3A/K4A formed infrequent assemblies that appeared much smaller and dimmer than WT foci, while R261A staining was diffuse without any indication of condensate formation (Figure 3E). Quantification of the number puncta per cell confirmed the abrogation of bright puncta formation in both mutants (Figure 3F). Flow cytometry experiments showed that ORF1p protein expression was similar across cells expressing the WT and mutant L1s, indicating that loss of puncta formation was likely due to defective assembly rather than decreased protein abundance (Figure 3-Supp 1A). These experiments revealed that ORF1p K3A/K4A and R261A are not able to form WT-like assemblies when expressed in the context of fulllength L1 in live cells, which contrasted with their mild condensation deficiencies in reconstitution experiments.

We wondered whether the loss of the ability to form bright ORF1p foci would have effects on L1 retrotransposition activity. A major advantage of our system is that these tagged ORF1p proteins were expressed in the context of an active L1 element with the GFP-AI retrotransposition reporter (Ostertag et al. 2000; An et al. 2011; Mita et al. 2018), which we used to assess the ability of each mutant to complete the entire L1 life-cycle. Using a well-characterized retrotransposition paradigm that involves 72 hours of L1 expression, we found that our tagged WT element retrotransposed in approximately 1% of cells. The K3A/K4A and R261A mutants, however, had undetectable retrotransposition activity (Figure 3G, Figure 3-Supp 1B). Taken together, these findings demonstrated that both K3/K4 and R261 are essential for ORF1p condensate formation in cells and retrotransposition, uncovering a possible connection between efficient ORF1p condensation and L1 retrotransposition activity.

### Dynamic coiled coils are essential for ORF1p condensation

Coiled coils are common structural motifs consisting of superhelical bundles of α-helices that promote multimerization of proteins. Coiled coil sequences are formed from characteristic heptad repeats, in which a pattern of hydrophobic and hydrophilic amino acids repeats every seven residues (Conway and Parry 1991). Insertions or deletions in this heptad repeat pattern have been shown to create local under- or overwinding of the supercoil that can affect coiled coil stability (Brown, Cohen, and Parry 1996). Previous work has characterized a three-residue stammer insertion (M91, E92, and L93) in the ORF1p coiled coil that, when deleted, increases the stability of the coiled-coil trimer and abrogates L1 retrotransposition (Khazina and Weichenrieder 2018). Since deletion of the stammer recovers the ideal heptad repeat pattern in the ORF1p coiled coil, we predicted that stammer-deleted ORF1p would maintain an elongated coiled coil conformation that might disfavor trimer-trimer interactions that are mediated by the N terminal half of the protein (Figure 4A, left two cartoons). As multivalency and dynamic interactions are key features of phase-separating proteins, we hypothesized that stammer-deleted ORF1p would be deficient in condensation due to a decrease in inter-trimer interactions. When we expressed L1 with a stammer-deleted (StammerDel) ORF1p in HeLa cells, we indeed found that the StammerDel protein was unable to form punctate condensates (Figure 4A, left two panels). We reasoned that there were two non-exclusive mechanisms by which stammer deletion led to the loss of ORF1p condensation: 1) the chemical properties of the stammer’s M, E, or L residues are necessary for condensation, or 2) a three-residue interruption in the heptad repeat pattern destabilized the coiled coil in a way that promoted condensation. To test these hypothetical mechanisms, we generated two additional ORF1p mutants with orthogonal stammers, one with the stammer residues mutated to three alanines (StammerAAA) and one with the wild-type E92 restored in the tri-alanine stammer (StammerAEA). We chose to specifically investigate the role of E92 because it was shown to be the only stammer residue that is conserved across primate L1s (Khazina and Weichenrieder 2018). Remarkably, both reconstituted stammer mutants formed puncta in cells (Figure 4A, right two panels). These experiments showed that destabilizing the coiled coil of ORF1p with a three-residue stammer insertion is necessary for condensate formation in cells.

**Figure 4:**
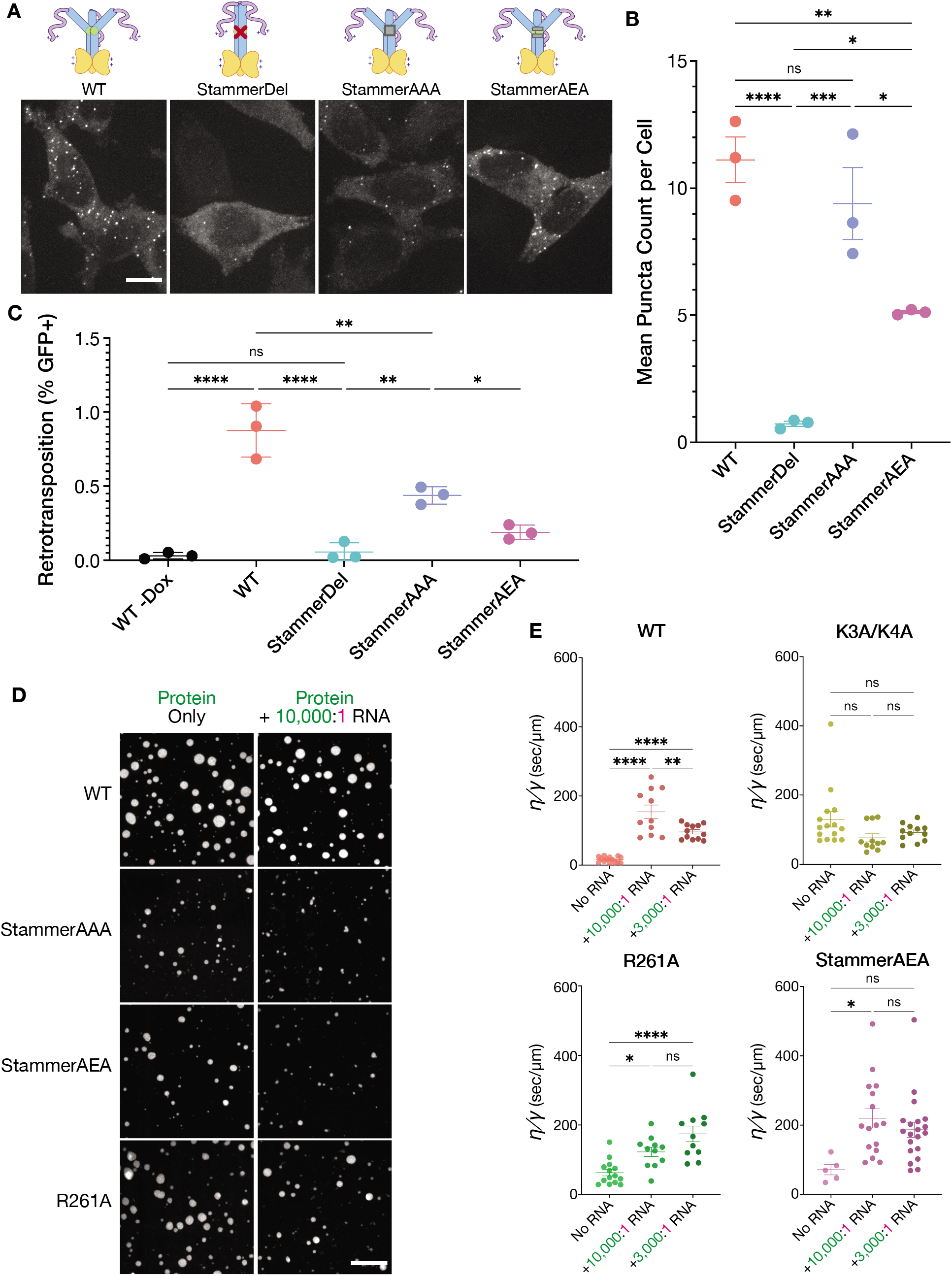
Stammer disruption of the ORF1p coiled coil is essential for L1 condensation in cells and modulates the physical properties of the ORF1p condensed phase. **(A)** Deletion of the stammer starkly decreases cellular ORF1p puncta formation, while stammer reconstitution rescues condensation. Representative maximum intensity Z projections of HeLa cells expressing WT, StammerDel, StammerAAA, or StammerAEA ORF1p after 6 hours of L1 expression. All images have the same lookup tables. Scale bar = 10 μm. **(B)** Stammer reconstitution rescues ORF1p puncta formation to WT-like levels. Quantification of the average number of ORF1p puncta per cell after 6 hours of expression of WT, StammerDel, StammerAAA, or StammerAEA ORF1p. Each point represents one biological replicate of induction and quantification and is the average of at least 75 cells. The mean and SEM of 3 biological replicates are shown, and statistical differences between mutants were calculated using a one-way ANOVA with Tukey’s multiple comparison correction. *p<0.05, **p<0.01, ***p<0.001, ****p<0.0001, ns = not significant. **(C)** ORF1p stammer deletion abrogates retrotransposition, while stammer reconstitution rescues retrotransposition activity. GFP+ cells were evaluated using FACS as in Figure 3G. The mean and SEM of 3 biological replicates are shown, and statistical differences between conditions were calculated using a one-way ANOVA with Tukey’s multiple comparison correction. *p<0.05, **p<0.01, ****p<0.0001, ns = not significant. **(D)** ORF1p with orthogonal stammer motifs form limited condensed phases that decrease in area in response to RNA addition. Representative images of the *in vitro* condensed phases of WT, StammerAAA, StammerAEA, and R261A ORF1p with and without the addition of 10000:1 RNA. Scale bar = 10 μm. **(E)** WT ORF1p and StammerAAA exhibit similar changes in inverse capillary velocity 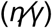 of their condensed phases in response to increasing RNA concentrations. Inverse capillary velocity was calculated from individual droplet fusion events in each condition, with each point representing a single analyzed fusion event; see methods for details. Mean ± SEM is shown; 5 or more fusion events were analyzed per condition. Changes in inverse capillary velocity are assessed across RNA conditions for each mutant independently using a one-way ANOVA with Tukey’s multiple comparison correction. *p<0.05, **p<0.01, ****p<0.0001, ns = not significant.

### Dynamic coiled coils that drive ORF1p condensation are essential for L1 retrotransposition

We next sought to better characterize the behaviors of the stammer mutants in cells and how they might affect retrotransposition. Quantifying the average number of puncta per cell confirmed the StammerDel variant’s inability to form condensates and revealed that StammerAAA formed a similar number of puncta per cell as WT, while StammerAEA’s puncta count was about half of that of WT (Figure 4B). We were surprised to find that restoring the wild-type E92 actually decreased ORF1p condensation; we noted that E92 is adjacent to a methionine residue in all primate ORF1p sequences (Khazina and Weichenrieder 2018), suggesting that the precise chemical properties of the stammer amino acids must be finely tuned for optimal activity. ORF1p expression in all three stammer mutant constructs was confirmed to be similar to WT expression, making it unlikely that these differences in condensation were due to protein abundance (Figure 4-Supp 1A). We then assayed the retrotransposition of these constructs and found that, while StammerDel was inactive, StammerAAA and StammerAEA both had readily detectable retrotransposition activity (50% for AAA, 20% for AEA relative to WT) (Figure 4C, Figure 4-Supp 1B). The pattern of retrotransposition activity strikingly mirrored the relative levels of puncta formation, with StammerAAA exhibiting a greater rescue of retrotransposition activity than StammerAEA, although both had less activity than WT. These findings demonstrated that synthetic stammer insertions are capable of driving ORF1p condensation and enable retrotransposition activity to an extent that correlates with the number of condensates formed per cell. Together, these experiments suggest that efficient ORF1p condensate formation is crucial for L1 retrotransposition.

### The physical properties of ORF1p-RNA condensates in vitro correspond with their ability to form puncta in cells

We purified full-length StammerAAA and StammerAEA ORF1p proteins with the aim of identifying *in vitro* condensed-phase behaviors that distinguish ORF1p variants that form puncta in cells from those that do not. Both stammer mutants were purified using the same protocol as WT and eluted as trimers indicating correct folding and assembly (Figure 4-Supp 2). We chose not to purify StammerDel as previous characterization of the stammer-deleted coiled coil alone required purification from inclusion bodies using denaturation followed by refolding (Khazina and Weichenrieder 2018), and it was unlikely that the full-length StammerDel ORF1p would refold in the correct conformation. When we assayed the AAA and AEA stammer mutants for condensation in buffer with physiological pH and salt concentration, we found that they formed round droplets but had a reduced total condensed-phase area compared to WT and even

R261A (Figure 4D, left). Adding low concentrations of labeled 2-kb L1 RNA to this condensation condition further reduced the condensate area of the stammer mutants, while the WT and R261A condensed phases were largely unaffected (Figure 4D, right). These experiments suggested that the formation of an extensive condensed phase *in vitro* is not a strong predictor of ORF1p condensation in cells.

Testing a wider range of RNA concentrations revealed that the stammer mutants are more sensitive to RNA than the other ORF1p variants (Figure 4-Supp 3A). Both stammer mutants exhibited further decreases in their already limited condensed phase areas with the addition of low concentrations of RNA (100,000:1 to 3,000:1 protein:RNA). Additionally, the stammer mutants both underwent extensive fibrillilarization at 1,000:1 protein:RNA, while WT and R261A retained a morphology similar to chains of slowly fusing droplets at the same RNA concentration. Similar to the stammer mutants, the ORF1p K3A/K4A condensed phase also underwent a fibrillar transition at 1,000:1 protein:RNA, despite being unaffected by lower RNA concentrations, suggesting that the N-terminal lysine residues play a role in keeping ORF1p soluble in the presence of RNA. These findings showed that the ORF1p stammer mutants, which robustly condense in cells, exhibit attenuated droplet formation compared WT ORF1p and the basic motif mutants in biochemical reconstitution experiments across a range of RNA concentrations.

Given the high sensitivity of the stammer-mutant condensates to RNA addition, we wondered whether the physical properties of their condensed phases, rather than their droplet areas, are the major predictors of their propensity to assemble in cells. We analyzed droplet fusion events for all mutants in the absence of RNA and in the presence of two RNA concentrations, 10,000:1 and 3,000:1 protein:RNA (Supp Movie 4). Notably, StammerAAA did not undergo a sufficient degree of condensation at either RNA concentration to evaluate fusion characteristics. Measuring the aspect ratio of droplet fusion events over time showed that in the absence of RNA, WT and StammerAEA exhibited faster fusion kinetics than the other mutants (Figure 4-Supp 3B, left). Given the extremely sharp reduction in aspect ratio over time for StammerAEA without RNA, it is likely that our difficulty in detecting fusion events for this mutant is a result of most fusions taking less than one minute to go to completion in this condition, preventing us from capturing them by imaging once per minute. However, with the addition of a low concentration of RNA (10,000:1 protein:RNA), both WT and StammerAEA exhibited much slower fusion kinetics than they did in the absence of RNA, while the behavior of other mutants was largely unchanged (Figure 4-Supp 3B, middle). At a higher RNA concentration, the fusion kinetics of all mutants were similar with the exception of R261A, which fused more slowly than the rest (Figure 4-Supp 3B, right).

Importantly, the kinetics of droplet fusion depend on both the sizes of the droplets and their physical properties. The characteristic fusion time *⊺* of two simple Newtonian liquid droplets suspended in a lower viscosity solution is given by 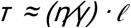, where *η* is the droplet viscosity, *γ* is the surface tension, and *ℓ* is the characteristic length scale, or size, of the droplets (Eggers, Lister, and Stone 1999; Brangwynne, Mitchison, and Hyman 2011).

The aspect ratio versus time plot of each analyzed fusion event was well fit by an exponential decay function, allowing us to extract a characteristic fusion time *⊺* for each fusion. We then divided each fusion event’s *⊺* by its characteristic length *ℓ* to determine the ratio of the condensed phase’s viscosity to its surface tension 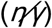, a value that is also known as its inverse capillary velocity (Brangwynne, Mitchison, and Hyman 2011). When we compared the inverse capillary velocity of WT ORF1p across the RNA conditions, we noted a complex effect that did not correlate with the amount of RNA added, as 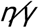 increased sharply with the addition of a low concentration of RNA and subsequently modestly decreased at a higher concentration of RNA (Figure 4E, top left). Interestingly, K3A/K4A and R261A had markedly different responses to RNA addition, with K3A/K4A showing no appreciable changes in inverse capillary velocity in response to RNA and R261A exhibiting increases in inverse capillary velocity with increasing RNA concentrations (Figure 4E, top right and bottom left). Significantly, StammerAEA’s inverse capillary velocity responded to RNA in a similar way to WT, exhibiting a steep increase at a low RNA concentration and a small decrease at the higher RNA concentration (Figure 4E, bottom right). Taken together, these experiments demonstrated that WT and StammerAEA ORF1p exhibit distinct non-linear changes in the inverse capillary velocity of their condensed phases in response to RNA, raising the possibility that differential condensed phase behavior in the presence of high and low RNA concentrations is important for condensate formation and L1 activity in cells.

## Discussion

Given the established importance of RNP formation for L1 retrotransposition and the recent appreciation of the contribution of dynamic interactions to the function of RNA-binding proteins, we investigated how the dynamic properties of ORF1p assemblies contribute to L1 retrotransposition. Although ORF1p’s major role in retrotransposition is thought to involve the formation of an RNP with the L1 mRNA and ORF2p in the cytoplasm, the diversity of ORF1p mutations that abrogate retrotransposition suggests the possibility of additional roles, including mediating RNP interactions with host proteins and participation in the downstream steps of nuclear translocation, DNA search for integration sites, or reverse transcription and integration (Sandra L. Martin et al. 2005; Kulpa and Moran 2005; Khazina et al. 2011; Taylor et al. 2013; Khazina and Weichenrieder 2018; Taylor et al. 2018; Adney et al. 2019). Using a well-characterized L1 expression system and modifying it to allow for live-cell imaging of ORF1p is a significant first step towards probing the behavior and function of ORF1p in cells. Expanding the use of this approach to investigate the role of ORF1p in cells and tissues with high endogenous L1 expression, including germline tissues and neoplastic tissues, will likely provide further insights into ORF1p interactions with host cell physiology (Branciforte and Martin 1994; Rodić et al. 2014; Ardeljan et al. 2017).

High spatiotemporal resolution imaging of ORF1p in live cells allowed us to investigate long-standing hypotheses regarding L1 RNP assembly. In particular, the ability of L1-encoded machinery to specifically drive retrotransposition of L1 mRNA while minimizing transposition of off-target substrates was hypothesized to occur through cotranslational binding of ORF1p and ORF2p to their encoding L1 mRNA in *cis* (Boeke 1997). This idea was substantiated with data demonstrating that two coexpressed L1 constructs exhibit only minimal *trans* complementation when one has a loss-of-function mutation in ORF1p or ORF2p (Wei et al. 2001; Kulpa and Moran 2006). Our work adds additional evidence supporting the *cis* preference model. The minimal mixing observed for co-expressed ORF1p puncta suggests that the physical properties of these assemblies minimize fusion events and protein exchange with the surrounding cytoplasm, providing a biophysical mechanism for the genetic *cis* preference (Figure 1F-G, Figure 4E). A mechanism involving reversible, gel-like co-condensation of ORF1p together with the RNA encoding it would allow for *cis* substrate preference without sequence-specific RNA binding (Kolosha and Martin 2003). We propose that ORF1p condensation in cells occurs cotranslationally, creating a viscous condensate with the L1 mRNA. Since this condensation must occur with the L1 mRNA from which it was translated for successful propagation, it requires a protein that is exquisitely sensitive to low concentrations of RNA, which we showed to be a biochemical characteristic shared by WT and StammerAEA (Figure 4E). Reduced inverse capillary velocities of WT and StammerAEA condensates at higher RNA concentrations may initially favor liquid-like growth of ORF1p condensates early in the RNP assembly process when protein:RNA ratios are lower, but as more ORF1p is generated and the protein:RNA ratio increases, our data suggest that the WT and StammerAEA condensates would become more viscous and gel-like in a way that K3A/K4A and R261A would not. This hypothetical role of ORF1p condensation in L1 RNP assembly would further imply that L1s expressing ORF1p defective in condensation may exhibit increased *trans* mobilization of off-target RNAs. Data supporting this conclusion has been reported for retrotransposition of the mammalian SINE *Alu,* which was observed to transpose twice as efficiently when driven by an L1 with ORF1p deleted rather than an intact L1 (Dewannieux, Esnault, and Heidmann 2003).

The behavior of ORF1p condensates in biochemical reconstitution experiments was distinct from the puncta properties observed in cells. In particular, the extensive droplet-like phase and the droplet fusion events seen in reconstituted ORF1p contrasted with the stereotyped punctate morphology and lack of mixing seen in cells. We found that reducing protein concentrations and titrating RNA concentrations allowed us to better approximate the punctate condensates seen in cells (Figure 2B). However, our reconstitution system did not fully recapitulate our observations in cells, in which WT ORF1p assembled but K3A/K4A and R261A did not (Figure 3B,F). Our simple biochemical reconstitution experiments involve equilibrium mixing of protein and RNA, allowing for nucleation, growth, and fusion of condensates in the absence of additional perturbations. While we posit that the discrepancies we observe are due to a cotranslational assembly mechanism in cells that is not reflected in our reconstituted system, there are many other buffer parameters and non-equilibrium processes in cells that could play a role in modifying ORF1p assembly kinetics, including macromolecular crowding, ATP hydrolysis, and post-translational modifications (Khan et al. 2018; Delarue et al. 2018; Brangwynne, Mitchison, and Hyman 2011; Brangwynne 2011; Nott et al. 2015; Aumiller and Keating 2016). Given this array of cellular parameters that may affect condensation of proteins in cells, we strove to ensure that we could characterize the behavior and properties of ORF1p assemblies in a cellular expression system that allowed us to simultaneously assess protein function. The ORF1 mutants in this study indicate that ORF1p condensation is required for L1 retrotransposition. We are now poised to further leverage this system to more precisely elucidate the emergent properties conferred to L1 RNPs as a result of condensation and how they contribute to ORF1p’s essential roles in retrotransposition.

A recent study also described phase separation behavior of ORF1p in biochemical reconstitution experiments (Newton et al. 2021). This study characterized the formation of an ORF1p condensed phase that is mediated by electrostatic interactions. The authors additionally demonstrated that a truncated ORF1p containing just the disordered NTR and coiled coil (residues 1-152; NTR-CC) was sufficient for phase separation. This description of NTR-CC sufficiency for phase separation in reconstitution experiments is in agreement with our characterization of R261A, which has greatly reduced structured RNA-binding activity (Khazina et al. 2011) but is still able to condense *in vitro.* We predict that both ORF1p R261A and the truncated NTR-CC protein phase-separate *in vitro* through a mechanism similar to complex coacervation, which requires only unstructured charge-charge interactions (Aumiller and Keating 2016; Boeynaems and Holehouse 2019). Our data suggest that the ORF1p RRM participates in protein-protein interactions that drive condensation in the absence of RNA, while the unstructured NTR plays a smaller role (Figure 3A-C). When ORF1p co-condenses with RNA, however, the NTR charge patch allows for modulation of the physical properties of the condensate in response to RNA (Figure 4E), which is critical for condensation in cells (Figure 3E-F). The RRM also contributes to the physical properties of ORF1p-RNA condensates, and we find that the combination of protein-protein and protein-RNA interactions of both the NTR basic patch and the RRM drives the formation of a condensate that is highly sensitive to low RNA concentrations (Figure 4E). Furthermore, the flexibility of the coiled coil that separates these two motifs appears to be critical to the roles of the NTR and RRM in condensation, since the StammerDel mutant that has a more rigid coiled coil exhibits starkly reduced condensation in cells (Figure 4A-B). In this way, we propose that two distinctive basic motifs and a dynamic coiled-coil scaffold allow ORF1p to undergo rapid condensation with *cis* RNA within the complex environment of the cell.

One important question that remains is whether ORF1p condensation is mechanistically required for L1 activity across species. Although aligning the ORF1 protein sequences across a diversity of species has proven difficult due to the variable nature of the N-terminal half of the protein, previous alignments have confirmed a shared ORF1p domain architecture across mammals that includes a likely trimeric coiled coil that varies in length, and RRM and CTD domains that allow for RNA binding (Boissinot and Sookdeo 2016; Khazina and Weichenrieder 2018). While the NTR is highly variable across mammals, the first eight residues of ORF1p contained at least two basic residues across a large diversity of mammals (Khazina and Weichenrieder 2018). Together, these sequence analyses suggest that diverse ORF1 proteins share the molecular features of multimerization, structured RNA binding, and unstructured electrostatic motifs that would enable them to undergo condensation. Studies of chimeric L1s have shown that substitution of the ORF1p in human L1 with that of a mouse L1 or an extinct megabat L1 generates elements that are still capable of undergoing retrotransposition in HeLa cells (Wagstaff, Barnerssoi, and Roy-Engel 2011; L. Yang et al. 2014). If ORF1p condensate formation is absolutely required for retrotransposition, these studies would predict that the mouse and megabat ORF1 proteins should undergo condensation in cells, which will be exciting to test in future studies.

The L1 system characterized in this work employs a uniquely powerful combination of biochemical reconstitution, live-cell imaging, and functional phenotyping in cells. *In vitro* reconstitution allows us to study the biophysical properties of condensates in a minimal and controllable system. We can use live-cell imaging to observe condensation in cells and identify factors and mutations that modulate condensate formation and behavior in the cellular milieu. A well-characterized retrotransposition assay enables us to correlate changes in cellular condensation with functional effects on the entire retrotransposon life-cycle. Further characterization of this system may reveal additional biophysical determinants of L1 retrotransposition, which will add to our understanding of functional biomolecular condensation within the complex cellular environment.

## Materials and Methods

### Plasmid construction

pLH2035 (pCEP-puro pTRE-Tight full-length L1RP containing ORF1-GGGGS-HaloTag7 and a GFP-AI cassette in its 3’ UTR) was generated from a pCEP4 episomal plasmid vector in which the hygromycin resistance cassette was substituted with a puromycin resistance cassette and the CMV promoter was swapped with a pTRE-Tight Tet inducible promoter as previously described; an untagged full-length L1RP sequence was then ligated downstream of the promoter (Taylor et al. 2013). ORF1-HaloTag7 was made using Gibson assembly (Gibson et al. 2009) of a PCR DNA fragment encoding GGGGS-HaloTag7 from pPM285 (pCEP-puro-ORFeus ORF1-HaloTag7, a gift from Jef Boeke) and pCEP-puro pTRE-Tight L1RP (Mita et al. 2018). The GFP-AI cassette was then digested and purified from pEA79 and ligated into the BstZ17I site in the 3’ UTR of L1RP (Mita et al. 2018). Localized mutations in ORF1p (K3A/K4A, R261A, StammerDel, StammerAAA, and StammerAEA) were introduced into pLH2035 using overlapping primers containing the mutation of interest to generate two PCR products that had homology to one another as well as to either a 5’ NotI site or a 3’ AfeI site that could then be Gibson assembled into the digested pCEP-puro-L1RP backbone, an approach similar to MISO mutagenesis (Mitchell et al. 2013).

pLH2060 (pCEP-puro pTRE-Tight full-length L1RP containing ORF1-GGGGS-mNeonGreen2) was generated from a pCEP-puro pTRE-Tight untagged L1RP construct. Gibson assembly was used to insert a GGGGS at the 3’ end of ORF1 as well as an AscI site for fluorophore insertion. The mNeonGreen2 sequence was PCR amplified from pLenti6.2_mNeonGreen2 (a gift from Vanessa LaPointe; Addgene plasmid # 113727) and was inserted in the AscI tagging site using Gibson assembly.

pMT538 (pETM11-6xHis-TEV-hORF1p) was a generous gift from Martin Taylor and Kathleen Burns (Carter et al. 2020). Localized mutations in ORF1p (K3A/K4A, R261A, StammerAAA, and StammerAEA) were engineered using Gibson assembly of the digested pMT538 backbone with two overlapping PCR products containing the mutation of interest and homology to either the BamHI or XbaI sites, as previously described.

All bacterial transformations for molecular cloning were done in DH10B competent cells (Thermo Fisher Scientific, product number EC0113). Oligonucleotide primers for cloning were ordered from IDT unless otherwise specified. All constructs were verified by Sanger sequencing (Genewiz).

### Protein purification

Purification of full-length wild-type and mutant ORF1p was performed using a modified version of a previously described protocol (Carter et al. 2020). pETM11-6xHis-TEV-hORF1p constructs were transformed into BL21(DE3) competent *E. coli* cells (Sigma-Aldrich, product number 69450) using a standard bacterial transformation protocol and were selected on LB + kanamycin agar plates. A single colony was grown in a 20 mL overnight culture in LB+Kan (1 x LB with 50 μg/mL kanamycin) at 37°C. The next day the cultures were diluted 1:100 in LB+Kan and the 2L of culture were grown in a shaker until they reached OD 0.8. The cultures were then transferred to a 16°C shaker for 1 hour, after which they were induced with 100 μM IPTG (EMD Millipore, product number 420322) overnight (18 hours) at 16°C. The rest of the purification was done at 4°C unless otherwise specified. The induced cultures were pelleted by spinning at 5,000g for 10 minutes. The pelleted culture was resuspended in 40 mL of cold lysis buffer (50 mM HEPES + 500 mM NaCl + 25 mM Imidazole + 1 mM DTT + 1X EDTA-free protease inhibitor cocktail (Thermo Fisher Scientific, product number A32965) pH 8) and was lysed using sonication in an ice-water bath. Following sonication, the sample was incubated with 200 units of benzonase (Sigma-Aldrich, product number E1014) for 30 minutes. The crude lysate was cleared with two clearing spins at 14,000g for 10 minutes. The cleared lysate was then incubated with 3 mL of 1:1 Ni-NTA agarose slurry (Qiagen, product number 30210) for 1 hour. Following pelleting, the Ni-NTA resin was washed twice with 10 mL of cold wash buffer (20 mM HEPES + 500 mM NaCl + 25 mM imidazole + 10 mM MgCl_2_ + 0.1% Triton X-100 (Fisher Scientific, product number BP151-500) pH 8). An additional 5-mL wash was done using wash buffer containing 200 units of benzonase, 1 μg/mL RNase A (Thermo Fisher Scientific, product number EN0531), and 1 mM DTT for 3 hours. Three additional 10-mL washes were done with cold wash buffer, and then the His-tagged protein was eluted from the resin in 5 mL of cold elution buffer (20 mM HEPES + 500 mM NaCl + 500 mM imidazole + 10 mM MgCl2 + 1 mM DTT pH 8) for 30 minutes. The eluted protein was then incubated with purified 6x-His-TEV protease E106G (Cabrita et al. 2007) and 500 μM TCEP (Thermo Fisher Scientific, product number 77720) for 20-24 hours. The cleavage mixture was then concentrated down to 500 μL in an Amicon-15 concentrator (EMD Millipore, product number UFC9050) and dialyzed back into lysis buffer overnight using small-volume dialysis chambers (Thermo Fisher Scientific, product number 88401). The protein mixture was subsequently incubated with 200 μL of Ni-NTA agarose slurry for 30 minutes to remove TEV protease as well as any uncleaved protein. The supernatant from this incubation wash then injected onto a Superose 6 Increase 10/300 GL size exclusion chromatography column (Cytiva, product number 29091596) on an AKTA pure protein purification system (Cytiva) and was run with cold, degassed gel filtration buffer (20 mM HEPES + 500 mM KCl pH 7.4). Fixed volume fractions were pooled based on A280 and A260 UV absorbance and ORF1p protein concentration was determined using NanoDrop A280 absorbance using a calculated molecular extinction coefficient of 25,440 M^-1^cm^-1^. The purified protein was then concentrated to 300 μM using an Amicon-2 Ultra concentrator (Millipore Sigma, product number UFC2030). DTT was added to the concentrated protein to a final concentration of 1 mM, and the protein was aliquoted, flash frozen in liquid nitrogen, and stored at −80°C.

For fluorescent labeling of purified ORF1p protein, approximately 10% of the purified protein was set aside prior to aliquoting and storage. Fluorescent ester dyes (green: Thermo Fisher Scientific, product number A37570; red: Thermo Fisher Scientific, product number A20003) were reconstituted in 10 μL of anhydrous DMSO (Thermo Fisher Scientific, product number D12345). The ORF1p protein was diluted to 500 μL in labeling buffer (20 mM HEPES + 500 mM KCl pH 6.5) and 1 μL of reconstituted dye was added to the solution and was mixed well by vortexing. The labeling mixture was then incubated mixing in the dark for 1 hour at 4°C, after which it was set to dialyze into ORF-KCl buffer (20 mM HEPES + 500 mM KCl + 1 mM DTT pH 7.4) overnight at 4°C. The protein concentration and moles dye per mole protein were calculated from NanoDrop absorbance measurements per manufacturer instructions. The protein was then aliquoted, flash frozen, and stored at −80°C.

The final purified sample as well as purification intermediates were checked for purity on protein gels. Samples were banked in 4X LDS Sample Buffer (Thermo Fisher Scientific, product number NP0007) with 5 mM DTT. Samples were boiled for 10 minutes at 95°C, vortexed thoroughly, and centrifuged before loading onto a precast gel (Thermo Fisher Scientific, product number NP0329) and running with MOPS buffer (Thermo Fisher Scientific, product number NP0001). The gel was then stained using SimplyBlue SafeStain (Thermo Fisher Scientific, product number LC6060) per manufacturer guidelines, and gel images were acquired using the 700 nm channel of an Odyssey CLx scanner (LiCOR).

### In vitro transcription and RNA purification

A DNA template containing the T7 promoter and the L1RP 5’ UTR and ORF1 sequence was generated using PCR of pLH2035 with a forward primer containing the T7 promoter and a reverse primer with that bound near the 3’ end of ORF1. The 1970 bp DNA was then gel purified and 500 ng of DNA template was loaded into a 20-μL *in vitro* transcription reaction with 5,000 units of T7 RNA polymerase (New England BioLabs, product number M0251LVIAL), 1X RNAPol Reaction Buffer (New England BioLabs, product number B9012SVIAL), 0.5 mM NTPs (New England BioLabs, product number N0450L), and 0.25 mM fluorescently labeled UTP. The labeled UTPs used were ChromaTide Alexa Fluor 488-5-UTP (Thermo Fisher Scientific, product number C11403), Cy3-UTP (Cytiva, product number PA53026), and Cy5-UTP (Cytiva, product number PA55026). The reaction mixture was incubated at 37°C for 24 hours and subsequently underwent RNA clean-up using a column-based purification (Qiagen, product number 74104). The purified RNA was eluted in nuclease-free water (Qiagen, product number 120114) and was quantified using NanoDrop A260 absorbance as well as the Qubit RNA HS assay (Thermo Fisher Scientific, product number Q32852). RNA samples were diluted to 300 nM and were aliquoted, flash frozen, and stored at −80°C.

### In vitro phase separation assays

Prior to starting the phase separation assays, the wells of a 384-well glass-bottom plate (Cellvis, product number P384-1.5H-N) were blocked as described in (Keenen, Larson, and Narlikar 2018). Briefly, the wells were treated with 2% Hellmanex (Sigma-Aldrich Z805939) for 1 hour, followed by three washes with ddH_2_O. Wells were subsequently treated with 1 M sodium hydroxide for 30 minutes, washed three times with ddH_2_O, and then incubated with freshly dissolved 20 mg/mL PEG-silane (Sigma-Aldrich JKA3037) in 95% ethanol overnight at room temperature. The plate was parafilmed to prevent evaporation and was stored in the dark. The next day the PEG-silane solution was removed, the wells were washed three times with ddH_2_O, and the wells were allowed to dry prior to plating protein mixtures.

ORF1p protein aliquots were thawed quickly at room temperature and were subsequently stored on ice, while fluorescently labeled RNA aliquots were thawed on ice. ORF1p unlabeled protein and labeled protein were mixed to a final concentration of 1-10% labeled protein and was subsequently diluted further with ORF1-KCl-Mg buffer (20 mM HEPES + 500 mM KCl + 1 mM MgCl2 + 1 mM DTT pH 7.4), if necessary. The RNA sample was diluted with ORF1 No-Salt-Mg buffer (20 mM HEPES + 1 mM MgCl_2_ + 1 mM DTT pH 7.4), if necessary. 25-μL mixtures were made containing a mixture of ORF1-KCl-Mg buffer and ORF1 No-Salt-Mg buffer to adjust the final salt concentration of the mixture. The protein was always the final component to be added to the mixture, and after its addition, the entire mixture was pipet mixed three times and plated immediately in a blocked well at room temperature. For end-point assays, the plated reactions were stored at room temperature in the dark for 2 hours prior to imaging. Fusion movies were imaged immediately following plating of all conditions, with images taken every minute. The plate was imaged on an Andor Yokogawa CSU-X confocal spinning disc on a Nikon TI Eclipse microscope with 488 and 640 nm lasers (Coherent) used to excite the labeled protein and RNA, respectively. Brightfield and fluorescence images were recorded using a Prime 95B scMOS camera (Photometrics) with a 100x objective (Plan Apo 100x DIC, Nikon, oil, NA = 1.45, part number = MRD01905, pixel size: 0.09 μm).

End-point assay images were loaded in FIJI, and each field of view (FOV) was segmented into two regions (condensed phase and background) based on the 488 protein fluorescence signal, with the condensed phase intensity needing to be at least 2x background intensity. Using the Analyze Particles function with a minimum particle size of 0.5 μm^2^, this segmented image was then used to calculate total condensed phase area, mean condensed phase protein intensity, and mean condensed phase RNA intensity, as well as protein partition coefficient and RNA partition coefficient by taking the ratio between the mean protein or RNA intensity in the condensed phase and the mean protein or RNA intensity in the background area. Total droplet area and protein and RNA partition coefficients were plotted using R, filtering out partition coefficient values for conditions with less than 1% of the field of view occupied by condensed phase.

Droplet fusion movies were loaded in FIJI and underwent 3D drift correction. Individual droplet fusion events were extracted from each movie, with the criteria that the fusion had to occur primarily within the focal plane and was not affected by a third droplet during the 15-minute fusion duration. Each fusion movie underwent segmentation to isolate the condensed phase, as previously described, and then the Analyze Particles function was used to fit an ellipse to the fusing droplets in order to determine the aspect ratio and the major and minor axis lengths of an ellipse fit to the fusing droplets in each frame of the 15-minute fusion. The aspect ratio vs. time plot for each fusion event was fit to a One Phase Decay non-linear fit in Prism 9 (GraphPad) with the constraints K > 0 and Plateau > 1, allowing for the extraction of values for fusion time constant *⊺* from each fusion event. Only fits with R^2^ values greater than 0.95 were used for analysis. Average aspect ratio vs. time plots for the fusions from each mutant-RNA condition were generated in Prism 9. In order to calculate the inverse capillary velocity, or the ratio of viscosity to surface tension 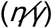, we used the equation 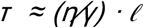, dividing the fusion time constant *⊺* by geometric mean ℓ = |(ℓ_major_(t = 0) – ℓ_minor_(t = 0)) · ℓ_minor_(t = 0)|^1/2^ (Eggers, Lister, and Stone 1999; Brangwynne, Mitchison, and Hyman 2011). The inverse capillary velocity for each analyzed fusion event was then plotted per mutant per RNA condition and differences in the mean and distribution of values was compared across RNA conditions for each mutant using a one-way ANOVA with Tukey’s multiple comparison correction.

### Mammalian cell culture

HeLa M2 cells (a gift from Gerald Schumann, Paul-Ehrlich-Institute; (Hampf and Gossen 2007)) were cultured in DMEM containing high glucose and sodium pyruvate (Thermo Fisher Scientific, product number 10313-039) supplemented with 10% FBS (Gemini, product number 100-106), 2 mM L-glutamine (Thermo Fisher Scientific, product number 25030-081), and 100 units/mL penicillin-streptomycin (Thermo Fisher Scientific, product number 15-140-122) (complete medium).

For transfection, HeLa M2 cells were seeded in a 6-well plate (Fisher Scientific, product number 50-202-137) on the day before transfection and were transfected with 1 μg of plasmid DNA per well using FuGENE-HD reagent (Promega, product number E2312) per manufacturer guidelines. HeLa M2 cells transfected with the L1 expression vector on a pCEP-puro episomal plasmid vector were selected and maintained in complete medium supplemented with 1 μg/mL puromycin dihydrochloride (MedChem Express, product number HY-B1743A); the supplemented media with puromycin was sterile filtered with a 0.22 μm filtration device (Worldwide Medical Products, product number 51101007) prior to use (complete puromycin media). Transfected HeLa M2 cells were considered “quasi-stable” for experimental use after at least 10 days of culture in complete puromycin media (Mita et al. 2020).

Cells were regularly split in fresh medium upon reaching 80-90% confluency. Cell culture medium was changed at least every 3 days. All cells were routinely tested for mycoplasma by PCR screening of conditioned medium.

### Mammalian live-cell imaging, puncta counting, and particle tracking

Quasi-stable HeLa cells with episomal inducible L1 expression constructs were seeded on non-coated 6-well, 12-well, or 96-well glass-bottom plates (Cellvis, product numbers P06-1.5H-N, P12-1.5H-N, and P96-1.5H-N) 1-2 days prior to imaging. Prior to imaging, the cells were induced with 1 μg/mL doxycycline hyclate (Sigma-Aldrich D9891) added to the conditioned media for the stated number of hours. The induced cells were then stained with 100 nM HaloTag Ligand JF549 (Promega, product number GA1111) and 500 nM SiR-DNA (Cytoskeleton, product number CY-SC007) in conditioned media per manufacturer instructions for 30-60 minutes. The cells were washed one time with DPBS (Fisher Scientific, product number 14-190-250) and fresh complete puromycin media was added to the cells for imaging.

Live cells were imaged on an Andor Yokogawa CSU-X confocal spinning disc on a Nikon TI Eclipse microscope equipped with a stage top incubation system (Tokai Hit and World Precision Instruments) to maintain a temperature of 37°C and 5% CO_2_ in the recirculated air. The Halo and SiR-DNA signals were obtained using 561 and 640 nm lasers (Coherent), respectively, and fluorescence images were captured using a Prime 95B scMOS camera (Photometrics) with a 60x objective (CFI Apo 60x, Nikon, oil, NA = 1.49, part number = MRD01691, pixel size: 0.13 μm). 10-μm Z stacks of Halo and SiR-DNA fluorescence were taken for puncta counting, with 1-μm steps between frames. For puncta tracking, Halo signal was acquired at 10 frames-per-second (fps) for 10 seconds and a matched SiR-DNA image was acquired to differentiate nuclear and cytoplasmic puncta.

All live-cell images were analyzed in FIJI. For puncta enumeration, Z stacks were projected using maximum intensity and cells were manually outlined using Halo signal and were saved to the ROI Manager. Puncta in the Z-projected image were then identified using the Find Maxima function with a Prominence value of 75 and the number of maxima located within each cellular ROI was recorded. Cells with extremely high expression levels were excluded due to the inability to correctly identify puncta using the fixed prominence value, and mitotic cells were excluded since the Z stack did not include their full volume. Cells with zero maxima were included. The mean and SEM of the puncta count per cell for each induction condition or mutant was determined in Microsoft Excel (Microsoft). The average and SEM of the puncta count per cell for each induction condition or mutant was determined across 3 biological replicates using Prism 9 (GraphPad) and was plotted. The puncta count per cell values were compared across conditions or mutants using a one-way ANOVA with Tukey’s multiple comparison correction where shown.

Puncta tracking was performed with the Mosaic suite of FIJI, using the following typical parameters: radius = 2, cutoff = 0, 20% of fluorescence intensity (Per/Abs), a link range of 1, and a maximum displacement of 5 px, assuming Brownian dynamics. Mean-square displacement (MSD) was then calculated for every 2D trajectory, selecting trajectories with more than 10 time points to reduce tracking error from particles moving in and out of the focal plane. We then fitted the time-averaged MSD of each selected trajectory with linear time dependence based on the first 10 time intervals (1 s): MSD(*⊺*)*⊺* = 4D_eff_*⊺*, where *⊺* is the imaging time interval and D_eff_ is the effective diffusion coefficient with the unit of *μm*^2^/*s*, which ignores the effects of sub-diffusion and superdiffusion on individual tracks but allows for better comparison across different conditions. We used the median value of D_eff_ among all trajectories within each field of view as a single data point and determined the mean and distribution of D_eff_ values across all fields of view. This analysis was implemented in a Python software developed in our lab called GEMspa (https://github.com/liamholtlab/GEMspa/releases/tag/v0.11-beta). Image areas corresponding to nuclei which contained tracked particles were manually outlined and a mask was generated that contained only those areas. The mask was inverted to create a mask containing all non-nuclear tracks, and these masks were used to analyze nuclear tracks and cytoplasmic (non-nuclear) tracks separately. Particle intensities were calculated using the average pixel intensities for a 5×5 square around each particle’s x,y position and were averaged across the length of the trajectory to generate a particle intensity for each track.

### Colocalization assay, colocalization analysis, and fixed cell imaging

Quasi-stable HeLa M2 cells transfected with the pLH2035 construct were transfected with pLH2060 in 6-well plates using Fugene-HD transfection reagent as previously described. The transfection mixture was replaced with complete puromycin media after 24 hours of incubation with the cells. The transfected cells were then seeded on non-coated glass-bottom 6-well plates (Cellvis, product number P6-1.5H-N) for 2 days. The cells were induced with doxycycline for 5 hours as previously described, and were subsequently stained with 100 nM Halo Ligand JF549 (Promega, product number GA1111) and 100 nM Halo Ligand JF646 (Promega, product number GA1121) for 1 hour in conditioned media containing doxycycline. The cells were then washed once with DPBS and were fixed in freshly diluted 4% formalin (Sigma-Aldrich, product number HT5012) for 10 minutes at room temperature. The cells were subsequently washed once with a PBS-Glycine solution (1X PBS + 10 mM glycine + 0.02% sodium azide + 0.2% Triton X-100 (Fisher Scientific, product number BP151-500)) and twice with 1X PBS. The cells were stained with 1 μM Hoechst 33342 (Thermo Fisher Scientific 62249) for 1 hour at room temperature in the dark and were subsequently stored in PBS at 4°C in the dark until imaging. The fixed plates of cells were imaged on an Andor Yokogawa CSU-X confocal spinning disc on a Nikon TI Eclipse microscope at room temperature. The fluorescence signals were obtained using DAPI epifluorescence excitation and the 488, 561, and 640 nm lasers (Coherent), and images were captured using a Prime 95B scMOS camera (Photometrics) with a 100x objective (Plan Apo 100x DIC, Nikon, oil, NA = 1.45, part number = MRD01905, pixel size: 0.09 μm). 8-μm Z stacks of Halo and SiR-DNA fluorescence were taken for puncta counting, with 1-μm steps between frames.

The colocalization images were analyzed in FIJI. The Z stacks were projected using maximum intensity and underwent background subtraction. 100 pixel x 100 pixel regions of interest within dual-expressing cells containing similar numbers of Halo+ and mNeonGreen2+ puncta were analyzed for colocalization using the Coloc2 package in FIJI. Two correlation values were determined for each ROI, one for Halo549-mNG2 (Halo-mNG2) and one for Halo549-Halo646 (Halo-Halo). Pearson’s R value for pixels above the threshold regression calculated using Bisection were used as the colocalization statistic. The paired Halo-mNG2 and Halo-Halo colocalization statistics for each ROI were then plotted on a dot plot in Prism 9 (GraphPad), and significant differences between Halo-mNG2 and Halo-Halo colocalization was determined using a two-tailed paired t test.

For qualitative imaging of ORF1-Halo in fixed cells, quasi-stable HeLa cells with episomal inducible L1 expression constructs were seeded in triplicate in glass-bottom 96-well plates (Cellvis, product number P96-1.5H-N) in complete puromycin media containing 1 μg/mL doxycycline for 72 hours. Cells were then stained with 100 nM Halo Ligand JF549 for 30 minutes in conditioned media containing doxycycline and washed once with DPBS. The cells were fixed as described above, using 4% formalin with washes with PBS-Glycine solution and PBS. Cells were stained with 100 nM SiR-DNA (Cytoskeleton, product number CY-SC007) for 1 hour at room temperature and were stored in the dark at 4°C until imaging. The fixed plates of cells were imaged on an Andor Yokogawa CSU-X confocal spinning disc on a Nikon TI Eclipse microscope at room temperature. The fluorescence signals were obtained using the 561 and 640 nm lasers (Coherent), and images were captured using a Prime 95B scMOS camera (Photometrics) with a 40x air objective (Plan Fluor 40x DIC, Nikon, air, NA = 0.75, part number = MRH00401, pixel size: 0.275 μm). 5×5 large images were acquired with 20% overlap using the ND Acquisition feature NIS Elements software (Nikon) and were stitched together using the SiR-DNA channel.

### Retrotransposition assays and FACS

Retrotransposition assays were conducted as described previously using quasi-stable HeLa M2 cells maintaining a pCEP-puro L1RP element containing a GFP-AI cassette (Mita et al. 2018). Briefly, quasi-stable HeLa cells were seeded in three separate wells of a 6-well plate (Fisher Scientific, product number 50-202-137) for each L1 construct using complete puromycin media with 1 μg/mL doxycycline. After 72 hours, the cells were washed once with DPBS and were stained with 1 μm Hoechst 33342 (Thermo Fisher Scientific, product number 62249) for 30 minutes at 37°C. The cells were then lifted using 500 μL of TrypLE Express Enzyme (Gibco, product number 12604021), neutralized with 500 μL of DMEM, and pelleted in 1.5-mL microcentrifuge tubes. The cell pellets were resuspended in 300 μL of freshly prepared FACS buffer (DPBS + 0.5% FBS), filtered into roundbottom tubes with cell strainer caps (Fisherbrand, product number 352235), and stored on ice until run on the Sony SH800S flow cytometer. 25,000 cells per condition were analyzed for GFP and Hoechst signals, which were compensated before analysis. The cutoff of GFP+ cells was set according to cells cultured for 72 hours in complete puromycin media without doxycycline, and the average percentage of GFP+ cells across all three technical replicate wells was reported as a single biological replicate for a given L1 construct. The average GFP+ rates across 3 biological replicates for each L1 construct were plotted in Prism 9 (GraphPad) and were compared using a one-way ANOVA with Tukey’s multiple comparison correction. Distributions of GFP intensity across cell populations were plotted using FlowJo (BD Biosciences).

For FACS analysis of cellular ORF1-Halo expression levels across ORF1p mutants, 2 million quasi-stable HeLa M2 cells maintaining a pCEP-puro L1RP element were seeded in a 10-cm tissue culture dish (Corning, product number 430167) in complete puromycin media. The following day, L1 expression was induced in the cells for 6 hours using 1 μg/mL doxycycline in the conditioned media. For the last hour of induction, the cells were stained with 100 nM HaloTag Ligand JF646 (Promega, product number GA1121) and 1 μm Hoechst 33342 (Thermo Fisher Scientific, product number 62249) at 37°C. As above, the cells were then washed once with DPBS, lifted using TrypLE Express Enzyme, and pelleted in 15-mL conical tubes (Corning, product number 430052). The cell pellets were resuspended in 2 mL of FACS buffer, filtered into round-bottom tubes with cell strainer caps and stored on ice until run on the Sony SH800S flow cytometer. Far-red fluorescence intensity was analyzed for 50,000 cells per ORF1p variant. Distributions of Halo intensity across cell populations were plotted using FlowJo (BD Biosciences).

## Supporting information

Supplemental figures and legends for movies

## Plasmid availability

All plasmids will be deposited in AddGene.

## Acknowledgements

We thank Paolo Mita for piloting initial live-cell L1 imaging and for extensive guidance through the vast collection of L1 expression constructs. We thank Martin Taylor for guidance on the purification and handling of bacterially-expressed ORF1 protein. We thank members of the Holt lab and Boeke lab for insightful scientific discussion and critical reading of the manuscript. ORF1p cartoon representations were created with BioRender.com.

This work was funded by NIH R01 GM132447 and R37 CA240765, the Chan Zuckerberg Initiative, and the Air Force Office of Scientific Research (AFoSR) grant FA9550-21-1-3503 0091 to LJH and a subaward from NIH P01 AG051449 to JDB. SS was funded by the Vilcek MSTP Scholars Award and NIH Medical Scientist Research Service Awards T32GM007308 and T32GM136573.

## References

Adney, Emily M., Matthias T. Ochmann, Srinjoy Sil, David M. Truong, Paolo Mita, Xuya Wang, David J. Kahler, David Fenyö, Liam J. Holt, and Jef D. Boeke. 2019. “Comprehensive Scanning Mutagenesis of Human Retrotransposon LINE-1 Identifies Motifs Essential for Function.” Genetics 213 (4): 1401–14.

An, Wenfeng, Lixin Dai, Anna Maria Niewiadomska, Alper Yetil, Kathryn A. O’Donnell, Jeffrey S. Han, and Jef D. Boeke. 2011. “Characterization of a Synthetic Human LINE-1 Retrotransposon ORFeus-Hs.” Mobile DNA 2 (1): 2.

Ardeljan, Daniel, Martin S. Taylor, David T. Ting, and Kathleen H. Burns. 2017. “The Human Long Interspersed Element-1 Retrotransposon: An Emerging Biomarker of Neoplasia.” Clinical Chemistry 63 (4): 816–22.

Aumiller, William M., Jr, and Christine D. Keating. 2016. “Phosphorylation-Mediated RNA/peptide Complex Coacervation as a Model for Intracellular Liquid Organelles.” Nature Chemistry 8 (2): 129–37.

Boeke, J. D. 1997. “LINEs and Alus--the polyA Connection.” Nature Genetics.

Boeynaems, S., and A. S. Holehouse. 2019. “Spontaneous Driving Forces Give Rise to Protein-RNA Condensates with Coexisting Phases and Complex Material Properties.” Proceedings of the. https://www.pnas.org/content/116/16/7889.short.

Boija, Ann, Isaac A. Klein, Benjamin R. Sabari, Alessandra Dall’Agnese, Eliot L. Coffey, Alicia V. Zamudio, Charles H. Li, et al. 2018. “Transcription Factors Activate Genes through the Phase-Separation Capacity of Their Activation Domains.” Cell 175 (7): 1842–55.e16.

Boissinot, Stéphane, and Akash Sookdeo. 2016. “The Evolution of LINE-1 in Vertebrates.” Genome Biology and Evolution 8 (12): 3485–3507.

Branciforte, D., and S. L. Martin. 1994. “Developmental and Cell Type Specificity of LINE-1 Expression in Mouse Testis: Implications for Transposition.” Molecular and Cellular Biology 14 (4): 2584–92.

Brangwynne, Clifford P. 2011. “Soft Active Aggregates: Mechanics, Dynamics and Self-Assembly of Liquid-like Intracellular Protein Bodies.” Soft Matter 7 (7): 3052.

Brangwynne, Clifford P., Christian R. Eckmann, David S. Courson, Agata Rybarska, Carsten Hoege, Jöbin Gharakhani, Frank Jülicher, and Anthony A. Hyman. 2009. “Germline P Granules Are Liquid Droplets That Localize by Controlled Dissolution/condensation.” Science 324 (5935): 1729–32.

Brangwynne, Clifford P., Timothy J. Mitchison, and Anthony A. Hyman. 2011. “Active Liquid-like Behavior of Nucleoli Determines Their Size and Shape in Xenopus Laevis Oocytes.” Proceedings of the National Academy of Sciences of the United States of America 108 (11): 4334–39.

Brouha, Brook, Joshua Schustak, Richard M. Badge, Sheila Lutz-Prigge, Alexander H. Farley, John V. Moran, and Haig H. Kazazian Jr. 2003. “Hot L1s Account for the Bulk of Retrotransposition in the Human Population.” Proceedings of the National Academy of Sciences of the United States of America 100 (9): 5280–85.

Brown, J. H., C. Cohen, and D. A. Parry. 1996. “Heptad Breaks in Alpha-Helical Coiled Coils: Stutters and Stammers.” Proteins 26 (2): 134–45.

Burns, Kathleen H., and Jef D. Boeke. 2012. “Human Transposon Tectonics.” Cell 149 (4): 740–52.

Cabrita, Lisa D., Dimitri Gilis, Amy L. Robertson, Yves Dehouck, Marianne Rooman, and Stephen P. Bottomley. 2007. “Enhancing the Stability and Solubility of TEV Protease Using in Silico Design.” Protein Science: A Publication of the Protein Society 16 (11): 2360–67.

Carter, Victoria, John LaCava, Martin S. Taylor, Shu Ying Liang, Cecilia Mustelin, Kennedy C. Ukadike, Anders Bengtsson, Christian Lood, and Tomas Mustelin. 2020. “High Prevalence and Disease Correlation of Autoantibodies Against p40 Encoded by Long Interspersed Nuclear Elements in Systemic Lupus Erythematosus.” Arthritis & Rheumatology (Hoboken, N.J.) 72 (1): 89–99.

Conway, J. F., and D. A. Parry. 1991. “Three-Stranded Alpha-Fibrous Proteins: The Heptad Repeat and Its Implications for Structure.” International Journal of Biological Macromolecules 13 (1): 14–16.

Cook, Pamela R., Charles E. Jones, and Anthony V. Furano. 2015. “Phosphorylation of ORF1p Is Required for L1 Retrotransposition.” Proceedings of the National Academy of Sciences of the United States of America 112 (14): 4298–4303.

Cost, Gregory J., Qinghua Feng, Alain Jacquier, and Jef D. Boeke. 2002. “Human L1 Element Target-Primed Reverse Transcription in Vitro.” The EMBO Journal 21 (21): 5899–5910.

Delarue, M., G. P. Brittingham, S. Pfeffer, I. V. Surovtsev, S. Pinglay, K. J. Kennedy, M. Schaffer, et al. 2018. “mTORC1 Controls Phase Separation and the Biophysical Properties of the Cytoplasm by Tuning Crowding.” Cell 174 (2): 338–49.e20.

Dewannieux, Marie, Cécile Esnault, and Thierry Heidmann. 2003. “LINE-Mediated Retrotransposition of Marked Alu Sequences.” Nature Genetics. https://doi.org/10.1038/ng1223.

Dombroski, B. A., S. L. Mathias, E. Nanthakumar, A. F. Scott, and H. H. Kazazian Jr. 1991. “Isolation of an Active Human Transposable Element.” Science 254 (5039): 1805–8.

Doucet, Aurélien J., Eugénia Basyuk, and Nicolas Gilbert. 2016. “Cellular Localization of Engineered Human LINE-1 RNA and Proteins.” Methods in Molecular Biology 1400: 281–97.

Doucet, Aurélien J., Jeremy E. Wilusz, Tomoichiro Miyoshi, Ying Liu, and John V. Moran. 2015. “A 3’ Poly(A) Tract Is Required for LINE-1 Retrotransposition.” Molecular Cell 60 (5): 728–41.

Eggers, Jens, John R. Lister, and Howard A. Stone. 1999. “Coalescence of Liquid Drops.” Journal of Fluid Mechanics 401 (December): 293–310.

Elbaum-Garfinkle, Shana, Younghoon Kim, Krzysztof Szczepaniak, Carlos Chih-Hsiung Chen, Christian R. Eckmann, Sua Myong, and Clifford P. Brangwynne. 2015. “The Disordered P Granule Protein LAF-1 Drives Phase Separation into Droplets with Tunable Viscosity and Dynamics.” Proceedings of the National Academy of Sciences 112 (23): 7189–94.

Feng, Q., J. V. Moran, H. H. Kazazian Jr, and J. D. Boeke. 1996. “Human L1 Retrotransposon Encodes a Conserved Endonuclease Required for Retrotransposition.” Cell 87 (5): 905–16.

Feric, Marina, Nilesh Vaidya, Tyler S. Harmon, Diana M. Mitrea, Lian Zhu, Tiffany M. Richardson, Richard W. Kriwacki, Rohit V. Pappu, and Clifford P. Brangwynne. 2016. “Coexisting Liquid Phases Underlie Nucleolar Subcompartments.” Cell 165 (7): 1686–97.

Gibson, Daniel G., Lei Young, Ray-Yuan Chuang, J. Craig Venter, Clyde A. Hutchison 3rd, and Hamilton O. Smith. 2009. “Enzymatic Assembly of DNA Molecules up to Several Hundred Kilobases.” Nature Methods 6 (5): 343–45.

Goodier, John L., Ling E. Cheung, and Haig H. Kazazian Jr. 2012. “MOV10 RNA Helicase Is a Potent Inhibitor of Retrotransposition in Cells.” PLoS Genetics 8 (10): e1002941.

Goodier, John L., Eric M. Ostertag, Kurt A. Engleka, Maria C. Seleme, and Haig H. Kazazian Jr. 2004. “A Potential Role for the Nucleolus in L1 Retrotransposition.” Human Molecular Genetics 13 (10): 1041–48.

Goodier, John L., Lili Zhang, Melissa R. Vetter, and Haig H. Kazazian Jr. 2007. “LINE-1 ORF1 Protein Localizes in Stress Granules with Other RNA-Binding Proteins, Including Components of RNA Interference RNA-Induced Silencing Complex.” Molecular and Cellular Biology 27 (18): 6469–83.

Hampf, Mathias, and Manfred Gossen. 2007. “Promoter Crosstalk Effects on Gene Expression.” Journal of Molecular Biology 365 (4): 911–20.

Hohjoh, H., and M. F. Singer. 1996. “Cytoplasmic Ribonucleoprotein Complexes Containing Human LINE-1 Protein and RNA.” The EMBO Journal 15 (3): 630–39.

Holmes, S. E., B. A. Dombroski, C. M. Krebs, C. D. Boehm, and H. H. Kazazian Jr. 1994. “A New Retrotransposable Human L1 Element from the LRE2 Locus on Chromosome 1q Produces a Chimaeric Insertion.” Nature Genetics 7 (2): 143–48.

Hyman, Anthony A., Christoph A. Weber, and Frank Jülicher. 2014. “Liquid-Liquid Phase Separation in Biology.” Annual Review of Cell and Developmental Biology 30: 39–58.

Januszyk, Kurt, Patrick Wai-Lun Li, Valerie Villareal, Dan Branciforte, Haihong Wu, Yongming Xie, Juli Feigon, Joseph A. Loo, Sandra L. Martin, and Robert T. Clubb. 2007. “Identification and Solution Structure of a Highly Conserved C-Terminal Domain within ORF1p Required for Retrotransposition of Long Interspersed Nuclear Element-1.” The Journal of Biological Chemistry 282 (34): 24893–904.

Jawerth, Louise, Elisabeth Fischer-Friedrich, Suropriya Saha, Jie Wang, Titus Franzmann, Xiaojie Zhang, Jenny Sachweh, et al. 2020. “Protein Condensates as Aging Maxwell Fluids.” Science 370 (6522): 1317–23.

Jurka, J. 1997. “Sequence Patterns Indicate an Enzymatic Involvement in Integration of Mammalian Retroposons.” Proceedings of the National Academy of Sciences of the United States of America 94 (5): 1872–77.

Kazazian, Haig H., Jr, and John V. Moran. 2017. “Mobile DNA in Health and Disease.” The New England Journal of Medicine 377 (4): 361–70.

Kazazian, H. H., Jr, C. Wong, H. Youssoufian, A. F. Scott, D. G. Phillips, and S. E. Antonarakis. 1988. “Haemophilia A Resulting from de Novo Insertion of L1 Sequences Represents a Novel Mechanism for Mutation in Man.” Nature 332 (6160): 164–66.

Keenen, Madeline M., Adam G. Larson, and Geeta J. Narlikar. 2018. “Visualization and Quantitation of Phase-Separated Droplet Formation by Human HP1α.” Methods in Enzymology 611 (October): 51–66.

Khan, Tarique, Tejbir S. Kandola, Jianzheng Wu, Shriram Venkatesan, Ellen Ketter, Jeffrey J. Lange, Alejandro Rodríguez Gama, et al. 2018. “Quantifying Nucleation In Vivo Reveals the Physical Basis of Prion-like Phase Behavior.” Molecular Cell 71 (1): 155–68.e7.

Khazina, Elena, Vincent Truffault, Regina Büttner, Steffen Schmidt, Murray Coles, and Oliver Weichenrieder. 2011. “Trimeric Structure and Flexibility of the L1ORF1 Protein in Human L1 Retrotransposition.” Nature Structural & Molecular Biology 18 (9): 1006–14.

Khazina, Elena, and Oliver Weichenrieder. 2009. “Non-LTR Retrotransposons Encode Noncanonical RRM Domains in Their First Open Reading Frame.” Proceedings of the National Academy of Sciences of the United States of America 106 (3): 731–36.

Khazina, Elena, and Oliver Weichenrieder. 2018. “Human LINE-1 Retrotransposition Requires a Metastable Coiled Coil and a Positively Charged N-Terminus in L1ORF1p.” eLife 7 (March). https://doi.org/10.7554/eLife.34960.

Kolosha, Vladimir O., and Sandra L. Martin. 2003. “High-Affinity, Non-Sequence-Specific RNA Binding by the Open Reading Frame 1 (ORF1) Protein from Long Interspersed Nuclear Element 1 (LINE-1).” The Journal of Biological Chemistry 278 (10): 8112–17.

Kulpa, Deanna A., and John V. Moran. 2005. “Ribonucleoprotein Particle Formation Is Necessary but Not Sufficient for LINE-1 Retrotransposition.” Human Molecular Genetics 14 (21): 3237–48.

Kulpa, Deanna A., and John V. Moran. 2006. “Cis-Preferential LINE-1 Reverse Transcriptase Activity in Ribonucleoprotein Particles.” Nature Structural & Molecular Biology 13 (7): 655–60.

Lander, E. S., L. M. Linton, B. Birren, C. Nusbaum, M. C. Zody, J. Baldwin, K. Devon, et al. 2001. “Initial Sequencing and Analysis of the Human Genome.” Nature 409 (6822): 860–921.

Lee, Chih-Yung S., Andrea Putnam, Tu Lu, Shuaixin He, John Paul T. Ouyang, and Geraldine Seydoux. 2020. “Recruitment of mRNAs to P Granules by Condensation with Intrinsically-Disordered Proteins.” eLife 9 (January). https://doi.org/10.7554/eLife.52896.

Li, Xiaoyu, Jianyong Zhang, Rui Jia, Vicky Cheng, Xin Xu, Wentao Qiao, Fei Guo, Chen Liang, and Shan Cen. 2013. “The MOV10 Helicase Inhibits LINE-1 Mobility.” The Journal of Biological Chemistry 288 (29): 21148–60.

Maharana, Shovamayee, Jie Wang, Dimitrios K. Papadopoulos, Doris Richter, Andrey Pozniakovsky, Ina Poser, Marc Bickle, et al. 2018. “RNA Buffers the Phase Separation Behavior of Prion-like RNA Binding Proteins.” Science 360 (6391): 918–21.

Martin, Sandra L., Margareta Cruceanu, Dan Branciforte, Patrick Wai-Lun Li, Stanley C. Kwok, Robert S. Hodges, and Mark C. Williams. 2005. “LINE-1 Retrotransposition Requires the Nucleic Acid Chaperone Activity of the ORF1 Protein.” Journal of Molecular Biology 348 (3): 549–61.

Martin, S. L., and F. D. Bushman. 2001. “Nucleic Acid Chaperone Activity of the ORF1 Protein from the Mouse LINE-1 Retrotransposon.” Molecular and Cellular Biology 21 (2): 467–75.

Mathias, S. L., A. F. Scott, H. H. Kazazian Jr, J. D. Boeke, and A. Gabriel. 1991. “Reverse Transcriptase Encoded by a Human Transposable Element.” Science 254 (5039): 1808–10.

Mita, Paolo, Xiaoji Sun, David Fenyö, David J. Kahler, Donghui Li, Neta Agmon, Aleksandra Wudzinska, et al. 2020. “BRCA1 and S Phase DNA Repair Pathways Restrict LINE-1 Retrotransposition in Human Cells.” Nature Structural & Molecular Biology 27 (2): 179–91.

Mita, Paolo, Aleksandra Wudzinska, Xiaoji Sun, Joshua Andrade, Shruti Nayak, David J. Kahler, Sana Badri, et al. 2018. “LINE-1 Protein Localization and Functional Dynamics during the Cell Cycle.” eLife 7 (January). https://doi.org/10.7554/eLife.30058.

Mitchell, Leslie A., Yizhi Cai, Martin Taylor, Anne Marie Noronha, James Chuang, Lixin Dai, and Jef D. Boeke. 2013. “Multichange Isothermal Mutagenesis: A New Strategy for Multiple Site-Directed Mutations in Plasmid DNA.” ACS Synthetic Biology 2 (8): 473–77.

Molliex, Amandine, Jamshid Temirov, Jihun Lee, Maura Coughlin, Anderson P. Kanagaraj, Hong Joo Kim, Tanja Mittag, and J. Paul Taylor. 2015. “Phase Separation by Low Complexity Domains Promotes Stress Granule Assembly and Drives Pathological Fibrillization.” Cell 163 (1): 123–33.

Moran, J. V., S. E. Holmes, T. P. Naas, R. J. DeBerardinis, J. D. Boeke, and H. H. Kazazian Jr. 1996. “High Frequency Retrotransposition in Cultured Mammalian Cells.” Cell 87 (5): 917–27.

Nanda, Jagpreet S., and Jon R. Lorsch. 2014. “Labeling a Protein with Fluorophores Using NHS Ester Derivitization.” Methods in Enzymology 536: 87–94.

Newton, Jocelyn C., Mandar T. Naik, Grace Y. Li, Eileen L. Murphy, Nicolas L. Fawzi, John M. Sedivy, and Gerwald Jogl. 2021. “Phase Separation of the LINE-1 ORF1 Protein Is Mediated by the N-Terminus and Coiled-Coil Domain.” Biophysical Journal 120 (11): 2181–91.

Nott, Timothy J., Evangelia Petsalaki, Patrick Farber, Dylan Jervis, Eden Fussner, Anne Plochowietz, Timothy D. Craggs, et al. 2015. “Phase Transition of a Disordered Nuage Protein Generates Environmentally Responsive Membraneless Organelles.” Molecular Cell 57 (5): 936–47.

Ostertag, E. M., E. T. Prak, R. J. DeBerardinis, J. V. Moran, and H. H. Kazazian Jr. 2000. “Determination of L1 Retrotransposition Kinetics in Cultured Cells.” Nucleic Acids Research 28 (6): 1418–23.

Pereira, Gavin C., Laura Sanchez, Paul M. Schaughency, Alejandro Rubio-Roldán, Jungbin A. Choi, Evarist Planet, Ranjan Batra, et al. 2018. “Properties of LINE-1 Proteins and Repeat Element Expression in the Context of Amyotrophic Lateral Sclerosis.” Mobile DNA 9 (December): 35.

Rodić, Nemanja, Reema Sharma, Rajni Sharma, John Zampella, Lixin Dai, Martin S. Taylor, Ralph H. Hruban, et al. 2014. “Long Interspersed Element-1 Protein Expression Is a Hallmark of Many Human Cancers.” The American Journal of Pathology 184 (5): 1280–86.

Sanders, David W., Nancy Kedersha, Daniel S. W. Lee, Amy R. Strom, Victoria Drake, Joshua A. Riback, Dan Bracha, et al. 2020. “Competing Protein-RNA Interaction Networks Control Multiphase Intracellular Organization.” Cell 181 (2): 306–24.e28.

Sharma, Reema, Nemanja Rodić, Kathleen H. Burns, and Martin S. Taylor. 2016. “Immunodetection of Human LINE-1 Expression in Cultured Cells and Human Tissues.” Methods in Molecular Biology 1400: 261–80.

Speek, M. 2001. “Antisense Promoter of Human L1 Retrotransposon Drives Transcription of Adjacent Cellular Genes.” Molecular and Cellular Biology 21 (6): 1973–85.

Swergold, G. D. 1990. “Identification, Characterization, and Cell Specificity of a Human LINE-1 Promoter.” Molecular and Cellular Biology 10 (12): 6718–29.

Tauber, Devin, Gabriel Tauber, Anthony Khong, Briana Van Treeck, Jerry Pelletier, and Roy Parker. 2020. “Modulation of RNA Condensation by the DEAD-Box Protein eIF4A.” Cell 180 (3): 411–26.e16.

Taylor, Martin S., Ilya Altukhov, Kelly R. Molloy, Paolo Mita, Hua Jiang, Emily M. Adney, Aleksandra Wudzinska, et al. 2018. “Dissection of Affinity Captured LINE-1 Macromolecular Complexes.” eLife 7 (January). https://doi.org/10.7554/eLife.30094.

Taylor, Martin S., John LaCava, Paolo Mita, Kelly R. Molloy, Cheng Ran Lisa Huang, Donghui Li, Emily M. Adney, et al. 2013. “Affinity Proteomics Reveals Human Host Factors Implicated in Discrete Stages of LINE-1 Retrotransposition.” Cell 155 (5): 1034–48.

Tóth-Petróczy, Agnes, Istvan Simon, Monika Fuxreiter, and Yaakov Levy. 2009. “Disordered Tails of Homeodomains Facilitate DNA Recognition by Providing a Trade-off between Folding and Specific Binding.” Journal of the American Chemical Society 131 (42): 15084–85.

Wagstaff, Bradley J., Miriam Barnerssoi, and Astrid M. Roy-Engel. 2011. “Evolutionary Conservation of the Functional Modularity of Primate and Murine LINE-1 Elements.” PloS One 6 (5): e19672.

Wei, W., N. Gilbert, S. L. Ooi, J. F. Lawler, E. M. Ostertag, H. H. Kazazian, J. D. Boeke, and J. V. Moran. 2001. “Human L1 Retrotransposition: Cis Preference versus Trans Complementation.” Molecular and Cellular Biology 21 (4): 1429–39.

Wheeler, Joshua R., Tyler Matheny, Saumya Jain, Robert Abrisch, and Roy Parker. 2016. “Distinct Stages in Stress Granule Assembly and Disassembly.” eLife 5 (September): 875.

Woodruff, Jeffrey B., Beatriz Ferreira Gomes, Per O. Widlund, Julia Mahamid, Alf Honigmann, and Anthony A. Hyman. 2017. “The Centrosome Is a Selective Condensate That Nucleates Microtubules by Concentrating Tubulin.” Cell 169 (6): 1066–77.e10.

Woods-Samuels, P., C. Wong, S. L. Mathias, A. F. Scott, H. H. Kazazian Jr, and S. E. Antonarakis. 1989. “Characterization of a Nondeleterious L1 Insertion in an Intron of the Human Factor VIII Gene and Further Evidence of Open Reading Frames in Functional L1 Elements.” Genomics 4 (3): 290–96.

Yang, Lei, John Brunsfeld, Luann Scott, and Holly Wichman. 2014. “Reviving the Dead: History and Reactivation of an Extinct l1.” PLoS Genetics 10 (6): e1004395.

Yang, Peiguo, Cécile Mathieu, Regina-Maria Kolaitis, Peipei Zhang, James Messing, Ugur Yurtsever, Zemin Yang, et al. 2020. “G3BP1 Is a Tunable Switch That Triggers Phase Separation to Assemble Stress Granules.” Cell 181 (2): 325–45.e28.

Zhang, Huaiying, Shana Elbaum-Garfinkle, Erin M. Langdon, Nicole Taylor, Patricia Occhipinti, Andrew A. Bridges, Clifford P. Brangwynne, and Amy S. Gladfelter. 2015. “RNA Controls PolyQ Protein Phase Transitions.” Molecular Cell 60 (2): 220–30.

Zhang, Yaojun, Daniel S. W. Lee, Yigal Meir, Clifford P. Brangwynne, and Ned S. Wingreen. 2021. “Mechanical Frustration of Phase Separation in the Cell Nucleus by Chromatin.” Physical Review Letters 126 (25): 258102.

Zwicker, David, Markus Decker, Steffen Jaensch, Anthony A. Hyman, and Frank Jülicher. 2014. “Centrosomes Are Autocatalytic Droplets of Pericentriolar Material Organized by Centrioles.” Proceedings of the National Academy of Sciences of the United States of America 111 (26): E2636–45.

