## Supplemental figures and legends for movies for "Condensation of LINE-1 is required for retrotransposition"

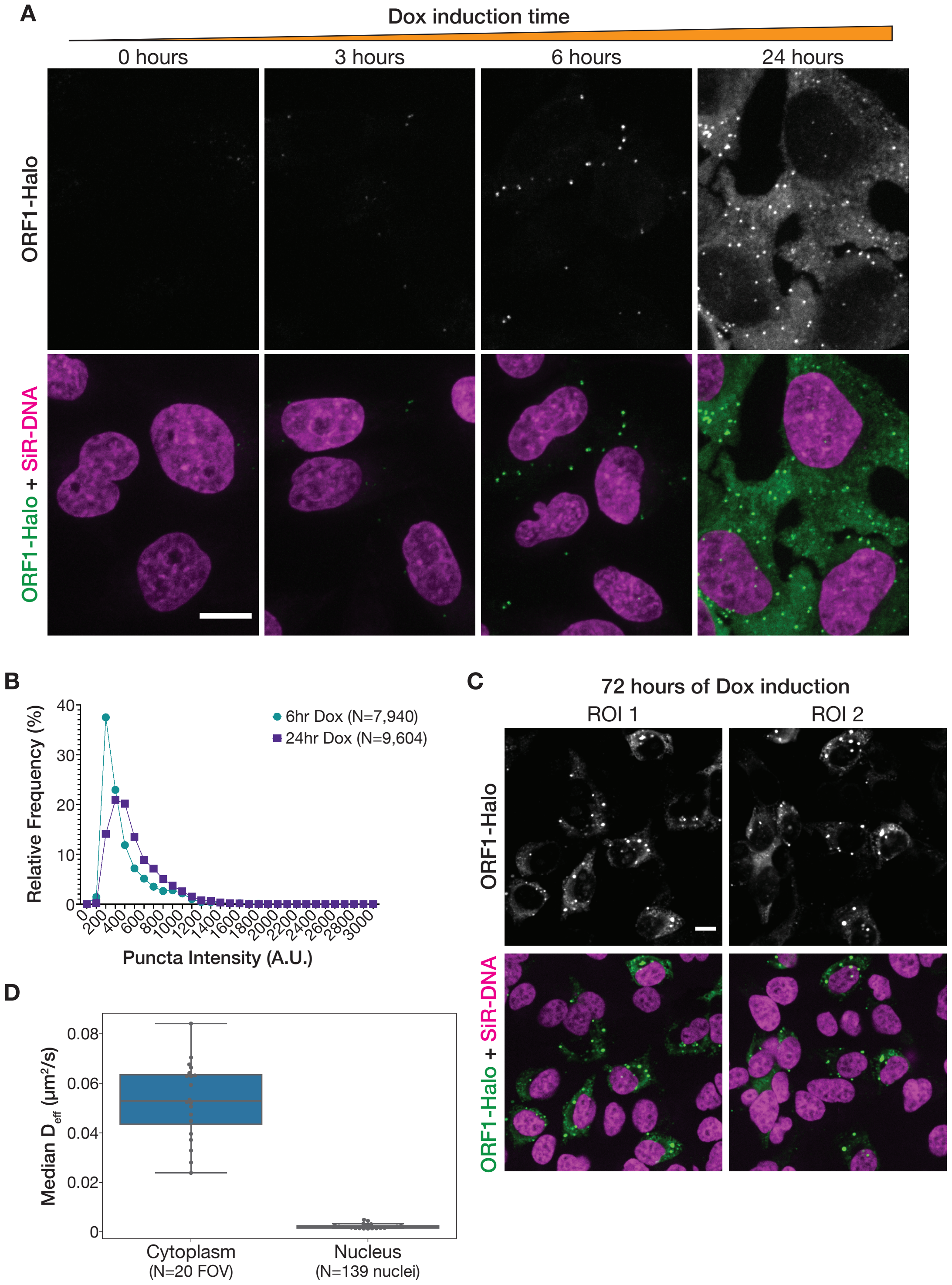


**Figure 1-Supplement 1: Increased L1 expression primarily increases the number of ORF1 puncta, but longer expression leads to the formation of larger stress-granule-like assemblies. (A)** ORF1 puncta form robustly after 6 hours of expression and increase in number after 24 hours of expression. Representative maximum intensity Z projections of live HeLa cells expressing WT ORF1p after varying lengths of L1 expression induction. ORF1-Halo signal is shown alone (top) and with nuclear staining (bottom, SiR-DNA). All of the ORF1p images have the same lookup tables. Scale bar = 10 μm. **(B)** The fluorescence intensity of ORF1 puncta increases slightly between 6 and 24 hours of L1 expression. Histograms of the intensities of ORF1 puncta in HeLa cells after 6 hours (N=7,940 puncta) or 24 hours (N=9,604 puncta) of L1 expression. Movies of ORF1 puncta were tracked and puncta intensities were averaged over the duration of each track (see methods for details). **(C)** Larger, more heterogeneous ORF1 condensates form after 72 hours of expression. Two representative ROIs of HeLa cells expressing L1 for 72 hours, stained with Halo ligand dye immediately prior to fixation. ORF1-Halo signal is shown alone (top) and with nuclear staining (bottom, SiR-DNA). Scale bar = 10 μm. **(D)** Nuclear ORF1 puncta diffuse much more slowly than their cytoplasmic counterparts. Movies of ORF1 puncta were tracked, and the median $D_{eff}$ for each nucleus (N=139) or each field of view excluding nuclei (FOV; N=20) was calculated and plotted as a single point (see methods for details). Boxplots show the spread of the median $D_{eff}$ values.

A

Transformation into BL21 bacterial competent cells

↓

Pre-culture overnight; inoculate large-scale culture

↓

Grow to OD ~0.8 at 37°C; induce with IPTG at 16°C overnight

↓

Sonication to lyse cells; benzonase treatment at 4°C ← Protein was kept in high-salt buffers (500 mM NaCl or KCl) and reducing conditions throughout the purification

↓

Two clear spins; binding incubation with Ni-NTA resin at 4°C

↓

Resin washes; longer on-bead RNase A and benzonase wash at 4°C

↓

Resin washes; elution; TEV protease cleavage of 6x-His tag overnight at 4°C

↓

Buffer exchange into low-imidazole buffer and run back on Ni-NTA resin, keeping the supernatant

↓

Size-exclusion chromatography (Akta); evaluate fractions, pool, and concentrate

↓

Amine-labeling using ester dyes; dialysis at 4°C

↓

Aliquot, flash freeze, and store in non-glycerol buffer at -80°C

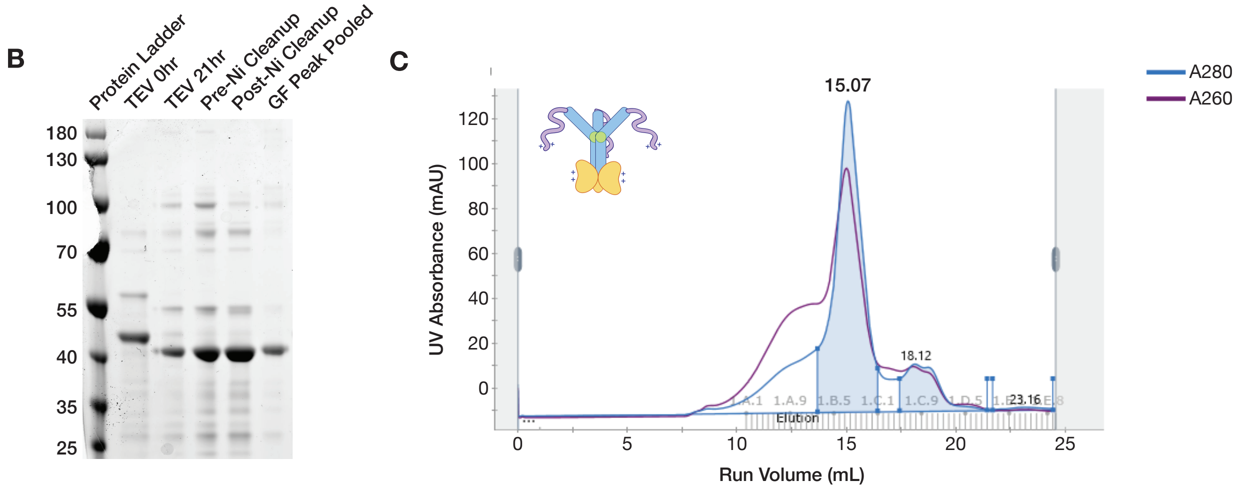


**Figure 2-Supplement 1: Full-length ORF1p purifies as a trimer from a bacterial expression system. (A)** A flowchart overview of the ORF1p purification protocol. A full purification protocol can be found in the methods. **(B)** Wild-type ORF1p can be purified with high purity from bacteria. A Coomassie gel shows purification intermediates from a purification of wild-type ORF1p. The gel filtration (GF) peak gel sample has a single dominant species at the expected size of 40 kDa. **(C)** Wild-type ORF1p expressed in bacteria purifies as a trimer. A representative UV absorbance trace of a GF run for wild-type ORF1p. The trace shows a single dominant protein species that elutes around 15 mL, corresponding to a molecular weight of 150 kDa per the column specifications. The blue shaded area under the primary peak indicates the fractions that were pooled for use in *in vitro* experiments. The A260 signal (purple) shows that the protein peak has a A260/A280 ratio of about 0.8, indicating low nucleic acid carryover. A large shoulder of higher molecular weight products that have much more nucleic acid and less protein can also be observed.

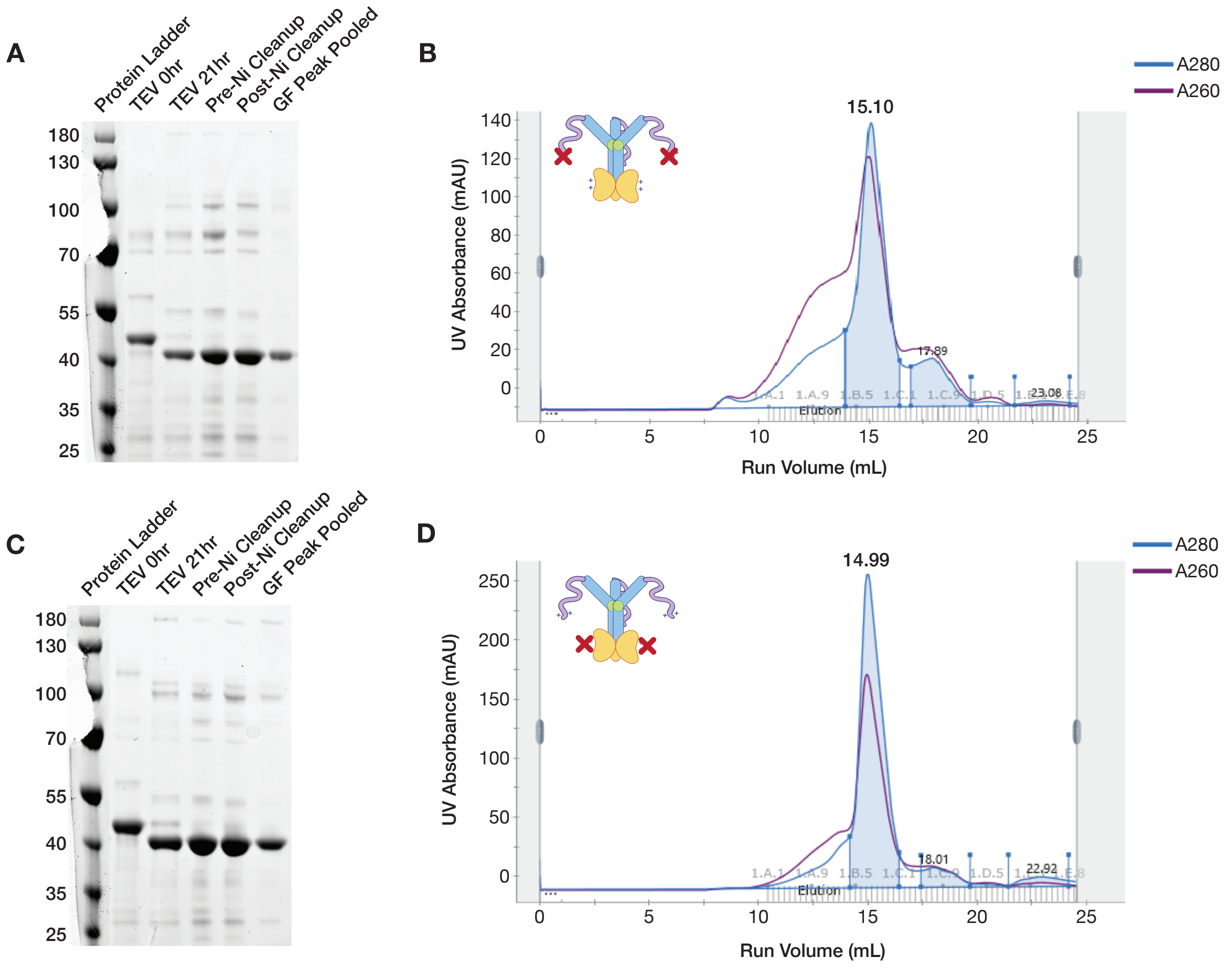


**Figure 3-Supplement 1: ORF1p K3A/K4A and R261A mutants purify as trimers. (A and C)** ORF1p K3A/K4A and R261A can be purified with high purity from bacteria. Coomassie gels showing purification intermediates from a purification of ORF1p K3A/K4A (A) and R261A (C), respectively. The gel filtration (GF) peak gel samples have a single dominant species at the expected size of 40 kDa. **(B and D)** ORF1p K3A/K4A and R261A expressed in bacteria purify as trimers. Representative UV absorbance traces of GF runs for ORF1p K3A/K4A (B) and R261A (D), respectively. The traces show a single dominant protein species that elutes around 15 mL, similar to WT. The blue shaded area under the primary peaks indicate the fractions that were pooled for use in *in vitro* experiments. The A260 signal (purple) shows that R261A has a much smaller shoulder of high-molecular-weight products with high nucleic acid carryover compared to WT or K3A/K4A, indicating that those species are driven by structured nucleic acid binding.

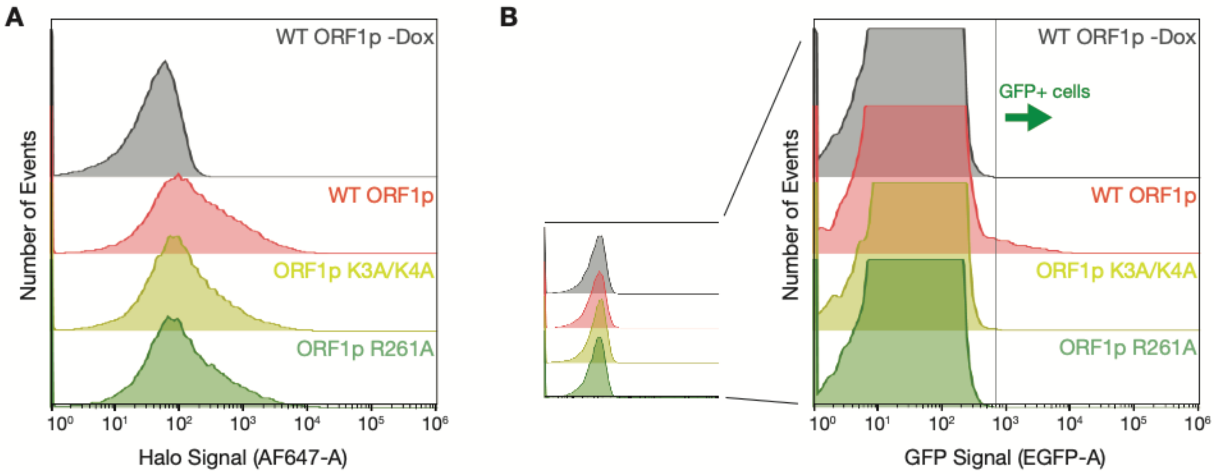


**Figure 3-Supplement 2: ORF1p basic mutants are expressed at similar levels to WT but have no retrotransposition activity. (A)** Cellular ORF1p expression levels are comparable across the basic motif variants following expression induction. FACS analysis of Halo signal from HeLa cells expressing WT, K3A/K4A, or R261A ORF1p for 6 hours, compared to WT ORF1p cells that did not undergo expression induction (WT ORF1p -Dox). 50,000 cells were analyzed per condition. **(B)** GFP+ cells from the GFP-AI retrotransposition reporter are detectable by FACS. Representative retrotransposition assay replicates from each condition are shown, exemplifying the clear detection of GFP+ cells in the induced WT ORF1p population but not in the other conditions.

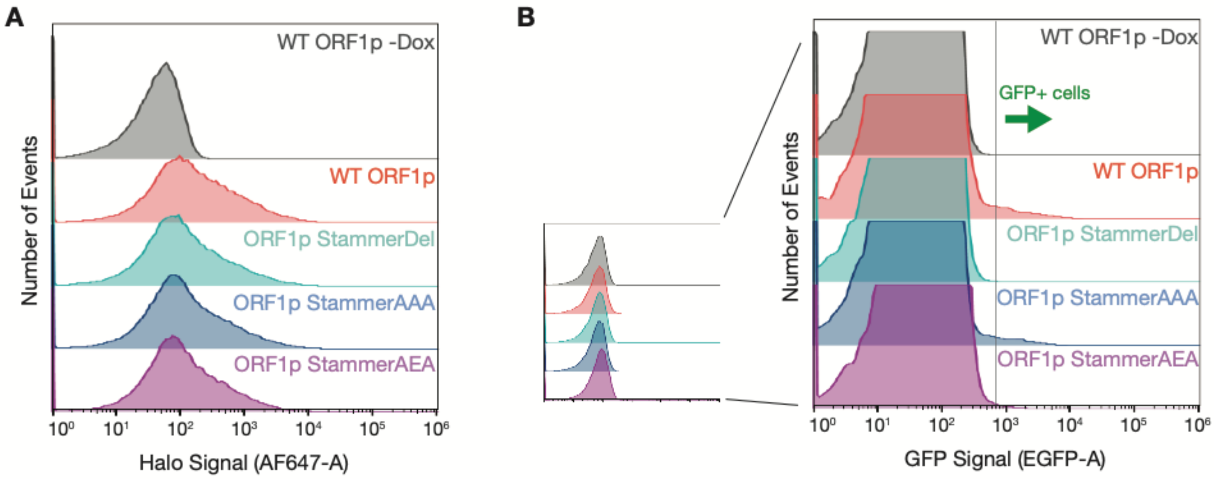


**Figure 4-Supplement 1: ORF1p stammer mutant proteins express at similar levels as WT but have variable retrotransposition rates. (A)** Cellular ORF1p expression levels are comparable across the stammer variants following expression induction. FACS analysis of Halo signal from HeLa cells expressing WT, StammerDel, StammerAAA, or StammerAEA ORF1p for 6 hours, compared to WT ORF1p cells that did not undergo expression induction (WT ORF1p -Dox). 50,000 cells were analyzed per condition. **(B)** GFP+ cells from the GFP-AI retrotransposition reporter are detectable by FACS. Representative retrotransposition assay replicates are shown, exemplifying the clear detection of GFP+ cells in the induced WT, StammerAAA, and StammerAEA populations.

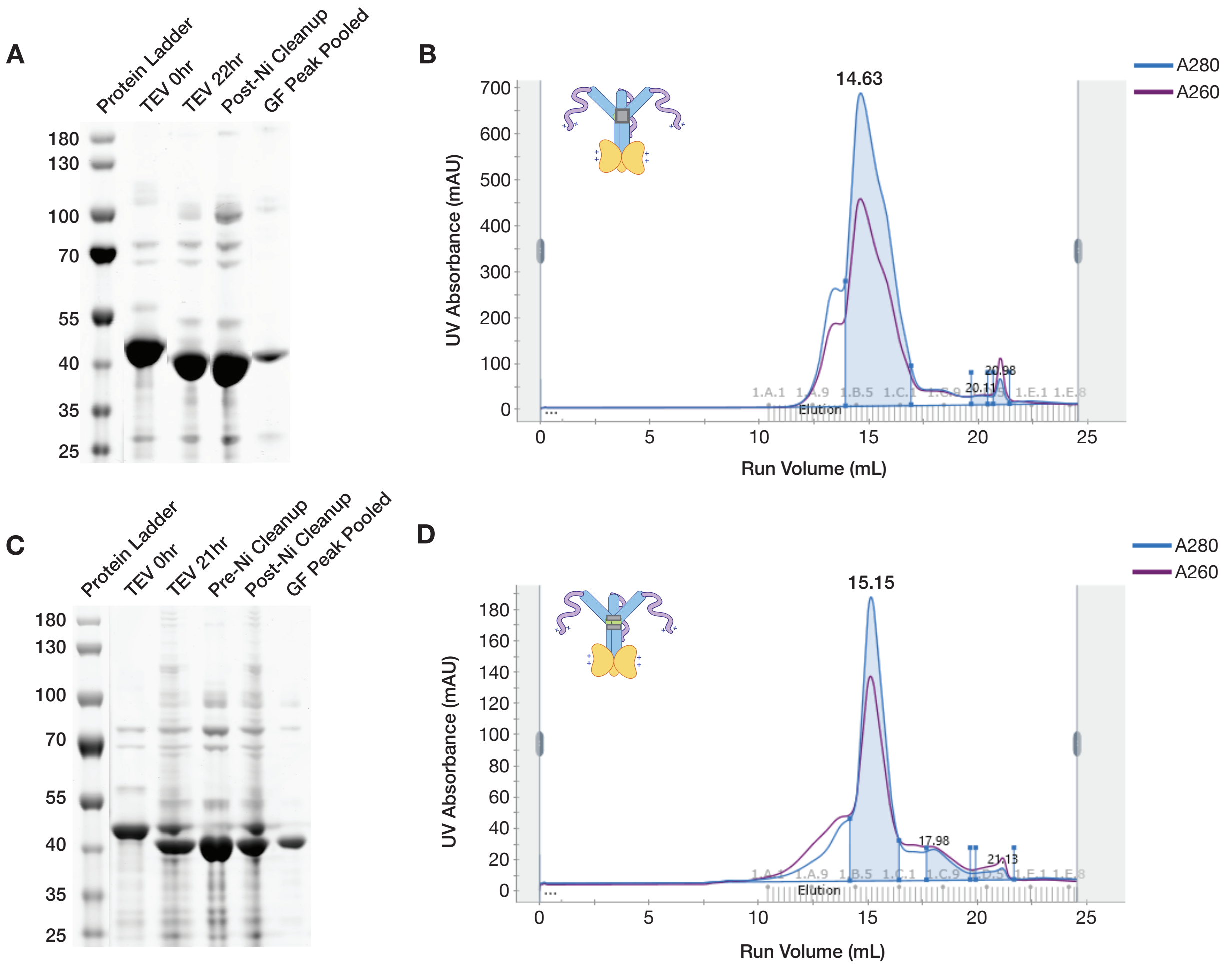


**Figure 4-Supplement 2: ORF1p StammerAAA and StammerAEA purify as trimers. (A and C)** ORF1p StammerAAA and StammerAEA can be purified with high purity from bacteria. Coomassie gels showing purification intermediates from a purification of ORF1p StammerAAA (A) and StammerAEA (C), respectively. The gel filtration (GF) peak gel samples have a single dominant species at the expected size of 40 kDa. **(B and D)** ORF1p StammerAAA and StammerAEA expressed in bacteria purify as trimers. Representative UV absorbance traces of GF runs for ORF1p StammerAAA (B) and StammerAEA (D), respectively. The traces show a single dominant protein species that elutes around 15 mL, similar to WT. The blue shaded area under the primary peaks indicate the fractions that were pooled for use in *in vitro* experiments. The A260 signal (purple) shows that both mutants have an A260/A280 of between 0.6 and 0.8, indicating low nucleic acid carryover.

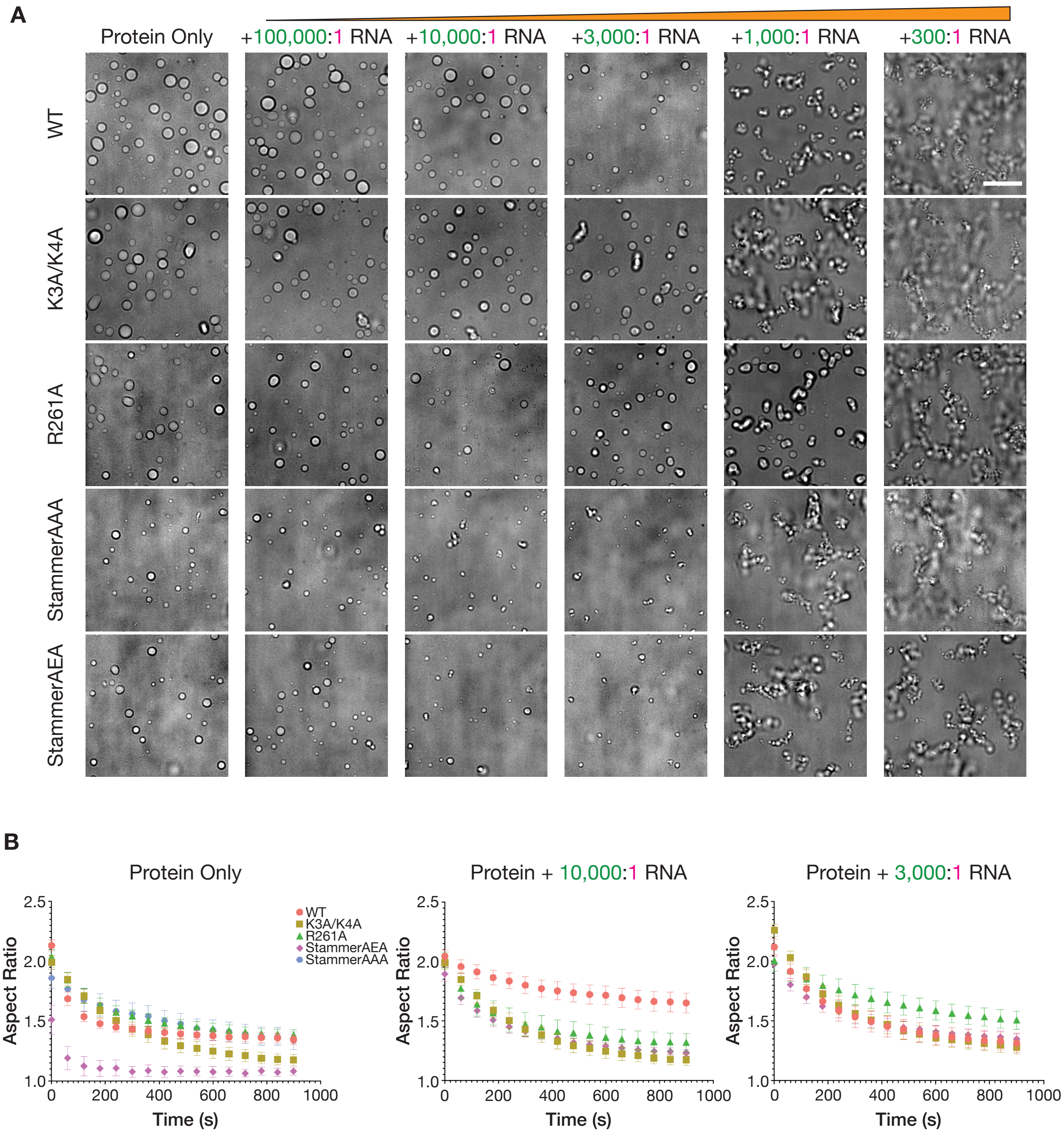


**Figure 4-Supplement 3: The physical properties of the condensed phases of ORF1p variants have differential responses to RNA. (A)** The StammerAAA and StammerAEA condensed phases are more sensitive to low RNA stoichiometries than the other mutants. Representative brightfield images of ORF1p mutant droplet morphology over a wide range of RNA stoichiometries. 5 µM protein and 150 mM KCl was used in all conditions. WT images are the same as in Figure 2B. Scale bar = 10 µm. **(B)** Fusion analysis of WT and mutant ORF1p droplets across RNA stoichiometries reveals differential fusion kinetics. 10 µM protein and 150 mM KCl were used in all conditions. Average aspect ratios across fusion events in each RNA condition (no RNA, 10,000:1

RNA, and 3,000:1 RNA) are plotted (Mean ± SEM) over time for 15 minutes. 5 or more fusions were analyzed per condition.

### Supplemental Movie Legends

**Supplemental Movie 1: ORF1 puncta diffuse more freely in the cytoplasm of HeLa cells than in the nucleus.** A representative real-time confocal microscopy movie of ORF1-Halo puncta diffusing in the cytoplasm and nuclei of HeLa cells after 6 hours of L1 expression (left), with a corresponding confocal image of nuclear staining with SiR-DNA (right). Scale bars = 10 μm.

**Supplemental Movie 2: Co-expressed ORF1 puncta exhibit minimal mixing in live HeLa cells.** A representative movie of a HeLa cell co-expressing two L1s with ORF1 tagged with either HaloTag7 or mNeonGreen2 (mNG2) for 4 hours. Confocal images in each channel were acquired every 5 seconds for 1 minute. Scale bar = 10 μm.

**Supplemental Movie 3: ORF1p droplets exhibit slower fusion kinetics in the presence of RNA.** Movies of ORF1p condensates settling out of solution and coalescing in the presence of varying amounts of RNA (from left to right: no RNA, +10,000:1 RNA, +3,000:1 RNA, and +1,000:1 RNA). 10 μM protein and 150 mM KCl were used in all conditions. Confocal images were acquired every minute for 2 hours. Scale bars = 5 μm.

**Supplemental Movie 4: Mutant ORF1p condensates exhibit differential changes in condensed phase properties with the addition of RNA.** Movies of ORF1p condensates settling out of solution and coalescing either in the presence of no RNA (top) or 10,000:1 RNA (bottom). All ORF1p variants (from left to right: WT, K3A/K4A, R261A, StammerAEA and StammerAAA) were assayed with 10 μM protein and 150 mM KCl. Confocal images were acquired every minute for 2 hours. Scale bars = 5 μm.
